# Rates and spectra of *de novo* structural mutation in *Chlamydomonas reinhardtii*

**DOI:** 10.1101/2022.05.23.493040

**Authors:** Eugenio López-Cortegano, Rory J. Craig, Jobran Chebib, Eniolaye J. Balogun, Peter D. Keightley

## Abstract

Genetic variation originates from several types of spontaneous mutation, including single nucleotide substitutions, short insertions and deletions (INDELs), and larger structural changes. Structural mutations (SMs) drive genome evolution and are thought to play major roles in evolutionary adaptation, speciation and genetic disease, including cancers. Sequencing of mutation accumulation (MA) lines has provided estimates of rates and spectra of single nucleotide and INDEL mutations in many species, yet the rate of new SMs is largely unknown. Here, we use long-read sequencing to determine the full mutation spectrum in MA lines derived from two strains (CC-1952 and CC-2931) of the green alga *Chlamydomonas reinhardtii*. The SM rate is highly variable between strains and MA lines, and SMs represent a substantial proportion of all mutations in both strains (CC-1952 6%; CC-2931 12%). The SM spectra also differs considerably between the two strains, with almost all inversions and translocations occurring in CC-2931 MA lines. This variation is associated with heterogeneity in the number and type of active transposable elements (TEs), which comprise major proportions of SMs in both strains (CC-1952 22% and CC-2931 38% of SMs). In CC-2931, a Crypton and a previously undescribed type of DNA element caused 71% of chromosomal rearrangements, while in CC-1952 a *Dualen* LINE was associated with 87% of duplications. Other SMs, notably many large duplications in CC-2931, were likely products of various double-strand break repair pathways. Our results demonstrate that diverse types of SMs occur at substantial rates and support prominent roles for SMs and TEs in evolution.

## Introduction

Since the development of the Modern Synthesis in evolutionary biology, the existence of chromosomal changes visualised in cytogenetic studies led to the hypothesis that structural mutations (SMs) could be an important source of variation leading to evolutionary change by natural selection (Dobzhansky and Epling, 1948; McClintock 1950; Ohno 1970). All genetic variation has its origin in new mutations, and efforts to estimate the rate of mutations started early in the 20th century by analysis of mutation accumulation (MA) experiments, in which spontaneous mutations are allowed to accumulate in lines of small effective population size where natural selection is ineffective (Muller 1928; Bateman 1959; Mukai 1964). The advent of whole-genome sequencing technology led to the possibility of directly estimating the rate, spectra and distribution of mutations in the genome by sequencing MA lines, and later by the sequencing of parents and their offspring. Studies of these kinds have been carried out in many species (Halligan and Keightley 2009; Yoder and Tiley 2021), but the short-read sequencing technology applied reliably detects only single nucleotide mutations (SNMs) and short insertions and deletions (INDELs), and little is known about the rates at which SMs occur *de novo*.

SMs include larger insertions and deletions (often defined as those >50 bp), duplications, transposable element (TE) insertions and excisions, and chromosomal rearrangements such as inversions and translocations. Such large structural changes may be expected to have larger fitness effects than SNMs and INDELs, and the structural variation that arises from SMs has been implicated in many evolutionary phenomena, including adaptation and speciation. For example, duplications can increase the copy number of functional sequences, and gene duplication provides the raw material for gene family evolution via processes including neo- and subfunctionalization (Kuzmin et al. 2022). Inversions may result in recombination suppression, and the subsequent evolutionary divergence of ancestral and inverted haplotypes has been implicated in local adaptation, speciation and sex chromosome evolution (Kirkpatrick and Barton 2006; Kirkpatrick 2010). Inversions may also give rise to ‘supergenes’, which preserve the linkage of multiple co-adapted loci and can underlie complex phenotypes (Joron et al. 2011, Küpper et al. 2015). Translocations and other major rearrangements can similarly suppress recombination and may directly cause reproductive isolation (Faria and Navarro 2010; Potter et al. 2017), while deletions have also been linked to genomic differentiation during speciation (Zhang et al. 2022). In particular, appreciation for the diverse evolutionary roles of TEs is ever increasing. TE insertions in functional sequences can drive rapid phenotypic adaptation (van’t Hof et al. 2016) and TEs can contribute substantially to regulatory sequence over evolutionary timescales (Chuong et al. 2017; Zhao et al. 2018). TE activity has been linked to phenomena including genome size evolution (Gregory 2005), genomic ‘turnover’ (gain and loss of DNA) (Kapusta et al. 2017) and speciation (Ricci et al. 2018; Tusso et al. 2022). TEs may also mediate large deletions, duplications and chromosomal rearrangements (Gray 2000), either directly as a byproduct of their transposition machinery (e.g. as in the classic *Ac*/*Ds* system (Zhang et al. 2009)) or via non-allelic homologous recombination between interspersed TE copies (Konkel and Batzer 2010). Finally, aside from their evolutionary importance, SMs are generally associated with deleterious effects and have been implicated in several human diseases and cancers (Weischenfeldt et al. 2013; Inaki and Liu 2012). SMs are also likely to become increasingly relevant in applied fields such as breeding, since structural variation has been associated with commercially important traits (Jayakodi et al. 2020; Song et al. 2020).

Fully understanding the evolutionary importance of structural variation requires a thorough knowledge of the rates at which the various types of SMs occur. Recently, advances in long-read sequencing technology have led to substantial improvements in the ability to assemble near-complete eukaryotic genomes and stimulated the development of bioinformatic tools for the discovery of structural variation (Rhoads and Au 2015; Jain et al. 2018; Mahmoud et al. 2019; Miga et al. 2020; De Coster et al. 2021). These advances now enable the study of *de novo* SMs in MA lines. Here, we use Pacific Biosciences (PacBio) HiFi sequencing to characterise the rates and spectra of SMs in two divergent strains of the single celled green alga *Chlamydomonas reinhardtii*. MA lines were generated in a previous study (Morgan et al. 2014) and we have previously used Illumina sequencing to investigate the rates and spectra of SNMs and INDELs (Ness et al. 2015). *C. reinhardtii* is an excellent model for mutation research, since its relatively large genome (∼111 Mb) and short generation time (∼2.5 generations per day) enables the rapid accumulation of large numbers of new mutations in a short time, and the species has been used to explore diverse genomic properties of new mutations (Ness et al. 2012; Sung et al. 2012; Böndel et al. 2021; López-Cortegano et al. 2021). We find that the rates and spectra of SMs differ substantially between MA lines and strains, and that SMs can represent a substantial proportion of the overall mutation rate.

## Results

### Structural mutation detection

We performed PacBio HiFi or continuous long read (CLR) sequencing of MA lines of the *C. reinhardtii* strains CC-1952 (*N*=4 MA lines, all HiFi) and CC-2931 (*N*=8, six HiFi and two CLR). These geographically distinct strains were selected on the basis of their relatively low (CC-1952, overall *μ* = 4.05 x 10^-10^ per site per generation) and high (CC-2931, *μ* = 15.6 x 10^-10^) SNM and INDEL rates estimated in our MA experiment, where lines were maintained for ∼1,050 generations by single cell descent (Morgan et al. 2014; Ness et al. 2015). To perform strain-specific detection of SMs in the MA lines, we first produced near-complete reference assemblies for both ancestral strains. The 17 *C. reinhardtii* chromosomes were assembled into 50 and 39 contigs with N50s of 4.25 and 3.81 Mb, for CC-1952 and CC-2931, respectively (Fig. S1). We subsequently defined nearly 98% of each ∼111 Mb ancestor genome as ‘callable” (i.e. sites where SMs could be called with high confidence; see Methods and Fig. S2). This represents a substantial increase over the ∼71% obtained in our previous study using short-read technology (Ness et al. 2015). We sequenced MA lines at sufficient depth of coverage to produce highly contiguous assemblies (Fig. S1), enabling us to call SMs using three approaches: directly from MA line PacBio read alignments against the appropriate ancestral reference using Sniffles (Sedlazeck et al. 2017), from MA line PacBio assembly alignments against the reference using MUM&Co (O’Donnell and Fischer 2020), and from Cactus pan-genomes using vg (Garrison et al. 2018; Armstrong et al. 2020). Sniffles and MUM&Co were run individually for each MA line, while all MA lines from each strain were analysed collectively with vg from a single strain-specific Cactus alignment. We subsequently collated and manually curated all variant calls using a combination of read visualisation and mapping approaches (Fig. 1A).

**Figure 1.**
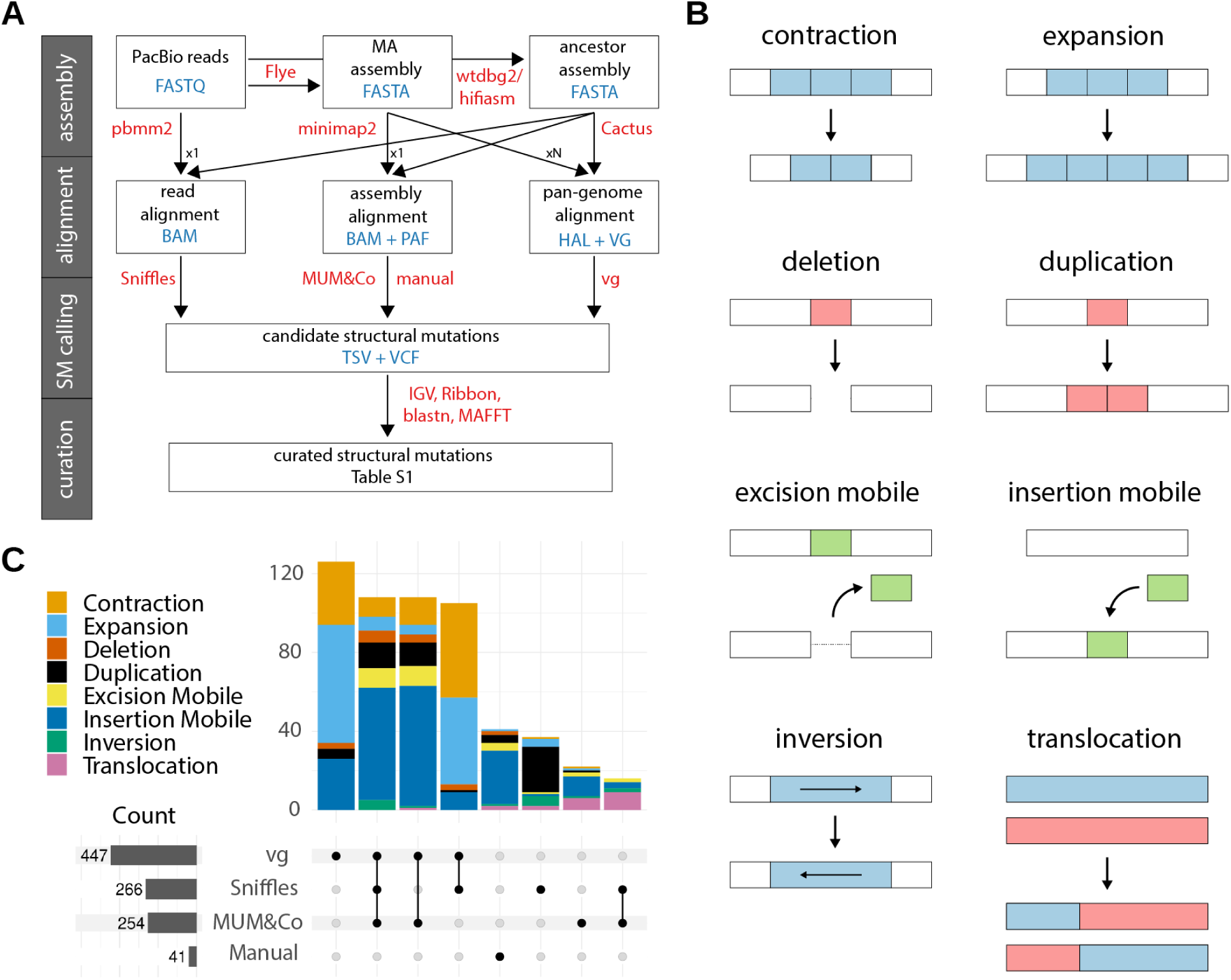
Structural mutation detection. A) Flowchart of the SM calling pipeline. Steps are organised from top to bottom in four stages: genome assembly, mapping and alignment, SM calling, and SM curation. Software used in each step is shown in red text, and file formats are in blue text. “Manual” indicates variants that were curated directly from alignment files. “x1” indicates that each dataset (reads or assemblies) was analysed individually, “xN” indicates that all MA lines for a given strain were analysed collectively. B) Schematics illustrating the eight different types of SM called. The ancestral state is shown above and the mutated state below. C) Intersection of the number of curated SMs identified by each calling method across all MA lines (for the two strains, CC-1952 and CC-2931, combined). In vertical bars, the number of SMs is coloured by SM type. Horizontal bars (in grey) show the total number of SMs called by each method.

We classified our curated dataset of SMs (a total of 120 in CC-1952 and 443 in CC-2931; Table S1) into eight categories: expansions and contractions of tandemly repeated sequence (e.g. in microsatellites or satellite DNA, collectively termed tandem repeat mutations, TRMs), duplications, deletions, insertions and excisions of mobile elements, and inversions and translocations (Fig. 1B). The different callers varied substantially in their ability to identify different SM types (Fig. 1C). Only 19.2% of SMs were called by all three tools, highlighting the importance of combining approaches. vg was most successful overall, calling 79.4% of SMs and 22.4% uniquely, although it called only 20.0% of inversions and translocations. MUM&Co called only 16.7% of TRMs, although it identified 64.5% of SMs in other categories. Sniffles called 47.2% of SMs and was generally outperformed by the assembly-based approaches, although it was superior in identifying duplications (calling 62.7% of duplications and 39.0% uniquely). The assembly-based methods failed to call duplications >30 kb (i.e. longer than the reads) since they were generally collapsed and absent in the assemblies. We also called 7.3% of SMs manually, most of which were transposable element (TE) insertions located at the breakpoints of other complex SMs (e.g. inversions and translocations).

The three approaches also differed markedly in the proportions of rejected calls (i.e. variants that could not be classified as genuine SMs), and all three returned more rejected calls than confirmed SMs (reaching 3x as many for vg; Fig. S3). Rejected calls generally fell in three categories: calls made within regions that we had defined as uncallable (which were not considered further), MUM&Co or vg calls that received assembly support but not read support (i.e. ‘valid’ calls introduced by assembly errors), or calls that received support from neither assemblies nor reads (unsupported variants). Considering the ratio of rejected calls to confirmed SMs in CC-2931, Sniffles performed best (1.14), followed by MUM&Co (1.81) and vg (3.42). However, these ratios varied considerably depending on the type and sequence context of the rejected call. All categories of rejected calls were associated with repetitive sequence (Fig. S4). The uncallable regions correspond to the most repetitive parts of the genome (generally very long satellite arrays), and despite representing only ∼2% of sites, a substantial proportion of calls were made in these regions (26.4%). Considering callable regions, tandem repeats of all lengths were a major source of both assembly errors and unsupported variants. After excluding all tandem repeats, the ratio of unsupported variants to confirmed SMs improved for all callers, falling from 0.21 to 0.17 for MUM&Co and from 0.91 to 0.20 for vg, although substantial proportions of calls attributed to assembly errors remained (Fig. S3). Consequently, although our results show that callers are capable of detecting SMs in tandemly repeated regions (expansions and contractions, particularly vg, Fig. 1C), attempting to do so runs the risk of introducing many false positives in fully automated pipelines. Rejected calls are further discussed in the Supplementary Material.

### Rates and spectra of structural mutations

The rates and spectra of SMs were remarkably different between the two strains (Fig. 2A, B). CC-2931 MA lines experienced ∼85% more SMs than CC-1952 MA lines, and total SM rates were significantly different between the strains (*μ*_SM (CC-1952)_ = 2.58 x 10^-10^ and *μ*_SM (CC-2931)_ = 4.30 x 10^-10^ per site per generation; W test, P = 4.04 x 10^-3^). However, the within-strain variance in *μ*_SM_ among MA lines was ∼12% higher than between them (ANOVA test, F = 10.84, P = 8.12 x 10 ^-3^). In terms of the number of bases affected, CC-2931 MA lines experienced larger SMs than CC-1952, and also experienced more types of SM (Fig. 2A). We observed only three SMs >20 kb in length in CC-1952 MA lines, whereas almost all large chromosomal rearrangements were found only in CC-2931, i.e. there were 1.75 inversions (median ∼243 kb) and 2.50 translocations per CC-2931 MA line, compared to a single 5.2 kb inversion across all CC-1952 MA lines. Deletions were also rare in CC-1952 (<1 per MA line on average), as were mobile excisions, due to a relative lack of active *cut-and-paste* DNA transposons (see below).

**Figure 2.**
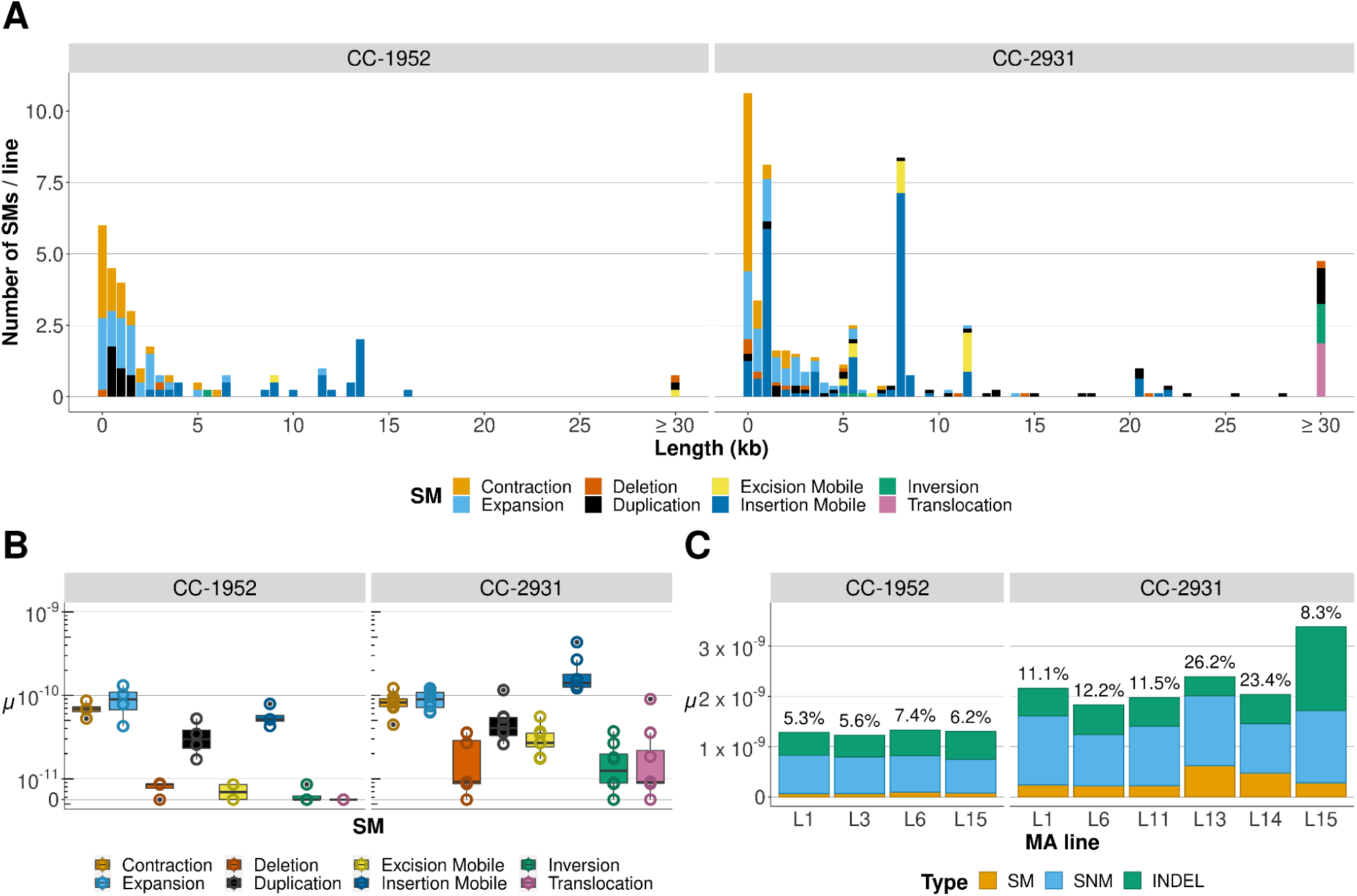
Spectrum and rate of structural mutations. A) Mean number of SMs by type (in colours) and length (in kb, rounded to 0.5 kb) per MA line. A total of 5 uncharacterized rearrangements of unknown length in CC-2931 MA lines are excluded. B) The SM rate per site per generation (*μ*, on a log_10_ scale) is plotted as open points and boxplots for different types of structural mutations. Data points represent individual MA lines. C) Mutation rates for different mutation types across the CC-1952 and CC-2931 MA lines. Only lines sequenced by PacBio HiFi are included. The percentage at the top of the bar indicates the proportion of SMs relative to all mutation types (SMs, SNMs and INDELs).

Although there were clear differences between the strains, some SM properties were shared by CC-1952 and CC-2931. TRMs were the most common category of SMs <3 kb in length in both strains (representing 60.8% and 35.0% of all SMs in CC-1952 and CC-2931, respectively) and occurred at similar rates between the two strains (median *μ*_TRM (CC-1952)_ = 1.68 x 10^-10^, *μ*_TRM (CC-2931)_ = 1.60 x 10^-10^, W test, P = 0.93). Mobile insertions dominated the spectra of SMs >3 kb in length (representing 21.7% and 37.9% of all SMs in CC-1952 and CC-2931, respectively), although the median rate of mobile insertions was ∼2.5x higher in CC-2931 (14.08 x 10^-11^) than in CC-1952 (5.15 x 10^-11^). Duplications were also relatively common in both strains (12.5% and 9.9% of all SMs in CC-1952 and CC-2931, respectively), although duplications in CC-1952 (median 0.9 kb) were significantly shorter than in CC-2931 (median 12.9 kb; W test, P = 2.98 x 10^-6^). Duplications were more frequent than deletions in the two strains (W test, P < 1.5 x 10^-2^), and also significantly longer in CC-2931 (median length 12.9 kb vs. 1.7 kb, respectively; W test, P = 1.81 x 10^-2^). Duplications and deletions affected coding and other genic sequences at rates similar to their genome-wide distributions (Fig. S5), suggesting that they could have considerable fitness effects.

In addition to identifying SMs, we used the PacBio HiFi reads to call SNMs and INDELs <50 bp in length. This analysis suggested that SMs represent 6.0% of the total mutation rate in CC-1952, and 11.7% in CC-2931. However, most mutations affecting short lengths of sequence were called in tandem repeats (∼60%), and given the uncertainty of variant calling in these sequences, we recalculated mutation rates excluding these regions. This had a minor effect on the relative contributions of SMs to the total rate (i.e. SMs represented 5.8% of mutations in CC-1952, and 11.8% in CC-2931; Fig. 2C). We did not find a significant correlation between *μ*_SM_ and either SNM or INDEL mutation rates among MA lines in either strain. Disregarding covariance terms between SM, SNM and INDEL mutation rates, variance in *μ*_SM_ explained 4.3% of total mutational variance in CC-1952 and 9.7% in CC-2931. Hence, our results suggest that SMs, including those that could have functional consequences, occur at a rate approximately 10-fold lower than the rate of SNMs and INDELs combined, yet their rates are highly variable between strains.

### Active transposable elements

With the exception of one putative mobile satellite in CC-2931 (see below), all mobile elements were TEs. We found that the two strains differed markedly in the number and diversity of active TE families. There were 12 families from seven TE subclasses active in CC-2931, and only three families from two subclasses active in CC-1952. We also observed considerable heterogeneity in insertion rates among MA lines and TE families (Fig. 2B, Fig. 3A). In CC-2931, per family rates ranged from a single insertion in only one MA line (e.g. the LINE *Dualen-4b_cRei*) to 62 insertions (minimum one per MA line, maximum 15) of the Crypton *CryptonF-1_cRei*.

**Figure 3.**
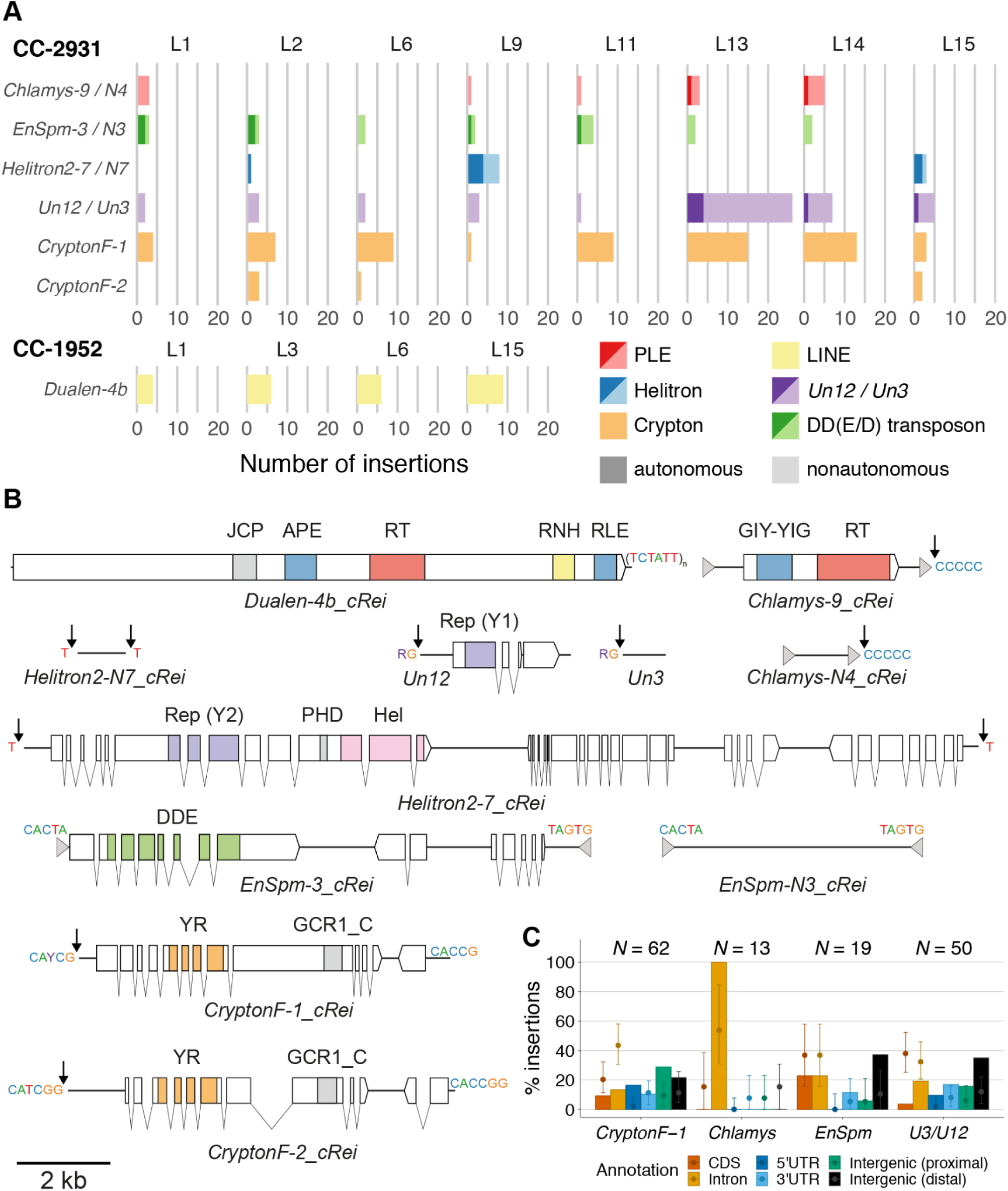
Active transposable elements. A) Number of insertions per TE family per MA line. Families with <2 insertions per strain are not shown. TE subclasses are shown in colours (red: PLE; yellow: LINE; blue: Helitron; purple: *Un12*/*Un3*; orange: Crypton; green: DD(E/D) transposon). Autonomous (darker colours) and nonautonomous (lighter colours) families that putatively rely on the same transposition machinery are grouped. The prefix of each TE name gives the superfamily (except *Un12/Un3*). B) Schematics of active TE families (to scale). Terminal inverted or direct repeats are shown by grey arrows, terminal sequences are shown above the main TE bodies (solid black lines), and insertion targets are shown next to black arrows. Coding sequence and introns of genes are shown by blocks and connecting lines, with domains coloured. RT = reverse transcriptase, GIY-YIG = GIY-YIG endonuclease, RNH = Ribonuclease H, APE = apurinic/apyrimidinic endonuclease-like endonuclease, RLE = restriction-like endonuclease, JCP = Josephin-related cysteine protease, Rep = replication protein (HUH endonuclease), Hel = helicase, PHD = plant homeodomain finger, DDE = DDE transposase, YR = tyrosine recombinase, GCR1_C = DNA-binding domain. C) Distribution of specific TE family insertions relative to genomic annotations in CC-2931. Autonomous and nonautonomous pairs were considered together. Error bars show a random expectation based on 1,000 insertions, adjusted for the genomic distribution of family-specific target sequences. Intergenic sequence is divided into proximal (within 500 bp of a gene) and more repetitive distal (>500 bp from a gene) sequences.

The most active retrotransposons in CC-2931 were an autonomous (*Chlamys-9_cRei*) and nonautonomous (*Chlamys-N4_cRei*) pair of *Penelope*-like elements (PLEs), which generally caused very short insertions (of median length 128 bp) due to 5’ truncation. All *Chlamys* insertions were into introns, which was broadly consistent with the underlying distribution of their (C)_n_ microsatellite target (Craig et al. 2021b), but was nonetheless more frequent than expected (Fig. 3B, C). The remaining CC-2931 TEs were various types of DNA transposons. The *cut-and-paste* DD(E/D) transposons *EnSpm-3_cRei* (autonomous) and *EnSpm-N3_cRei* (nonautonomous) were active in all lines. *EnSpm* insertions were significantly enriched in intergenic regions distant from genes, although unlike other TE families they were not underrepresented in coding sequence (Fig. 3C). Consistent with a higher efficiency of nonautonomous transposons relative to their longer autonomous counterparts (Han et al. 2013), we observed a net increase in copies of *EnSpm-N3_cRei* (three ancestral copies compared to a mean of 3.6 copies in the MA lines) and decrease in *EnSpm-3_cRei* (three ancestral copies compared to a mean of 2.4 copies in the MA lines). The net increase in *EnSpm-N3_cRei* copies can presumably be attributed to transposition during DNA replication (Feschotte and Pritham 2007). Also note that the number of insertions for these elements are minimum estimates (Fig. 3A), since several *cut-and-paste* transpositions may have occurred during the experiment that were not captured at the final time point.

We observed 12 insertions of *copy-and-paste* Helitrons, including the particularly long 20.4 kb autonomous *Helitron2-7_cRei* (Fig. 3B). We also observed an unusual pair of *copy-and-paste* transposons, here named *Un12* (autonomous) and *Un3* (nonautonomous), which inserted upstream of “RG” target sequences (R indicating purine) and caused variable length target site duplications. *Un12* contains a gene with introns that encodes a protein containing an HUH endonuclease of the Rep superfamily, a widespread domain in prokaryotes and viruses that is also found in eukaryotic Helitrons and prokaryotic IS*200*-IS*605* and IS*91*/ISCR transposons (Kazlauskas et al. 2019). However, since the *Un12* protein lacked a helicase domain and featured only one catalytic tyrosine (i.e. Y1 rather than Y2, Fig. 3B), these elements appear to be the first known members of a new group of eukaryotic TEs. We have since found similar TEs in several other taxa and their wider distribution will be described elsewhere. *Un3* was the second most active TE, with insertions in all lines and a maximum of 22 insertions in L13 (Fig. 3B). *Un3* and *Un12* insertions were significantly underrepresented within coding sequence and introns (Fig. 3C).

We observed two families of autonomous Cryptons, including *CryptonF-1_cRei,* which was the most active TE in CC-2931. To our knowledge these are the first observations of active Cryptons. Goodwin et al. (2003) proposed a model of insertion via site-specific recombination between a short donor sequence at one terminus of a Crypton and a near-identical target sequence at the integration site, catalysed by the Crypton-encoded tyrosine recombinase. Our data were consistent with this model, e.g. *CryptonF-1_cRei* terminated in the motif “CACCG” and targeted “CAYCG” (Y indicating pyrimidine; Fig. 3B). However, it was also proposed that Cryptons would undergo excision. While we observed clean excisions that left behind only the “CAYCG” target, the Cryptons increased in copy number, suggesting a more complex mode of transposition. *CryptonF-1_cRei* insertions were enriched at gene-proximal intergenic sequences and 5’ untranslated regions (UTRs).

In contrast to CC-2931, only the LINE retrotransposon *Dualen-4b_cRei* was active in multiple CC-1952 MA lines (with a range of four insertions in MA line 1 to nine in line 15, Fig. 3A). We also observed two *cut-and-paste* DD(E/D) transposons: a single excision and insertion of a *P* element (*P-2_cRei*) in MA line 1 (i.e. L1), and an excision (and extinction) of a giant single-copy 31.7 kb *Zisupton* DNA transposon (*Zisupton-3_cRei*) in L3.

### Transposable element-mediated structural mutations

We found that TE insertion and excision events were associated with many other SM types (Table 1). The number of TE insertions was positively correlated with the combined number of deletions, duplications, inversions and translocations across all MA lines (Pearson’s product-moment correlation, t_10_ = 2.64, *r* = 0.64, P = 0.02). Inversion and translocation breakpoints had a similar genomic distribution as TE insertions, exhibiting a bias towards UTRs and intergenic sequences (Fig. S5). More specifically, 70.6% of inversions and translocations in CC-2931 MA lines involved either *CryptonF-1_cRei* or *Un3/Un12* elements at one or both of the breakpoints (Table 1), with one exception that involved truncated copies of both *Un3* and *CryptonF-1_cRei*. We also attributed a far smaller number of SMs to homology-mediated double-strand break (DSB) repair mechanisms (Table 2), which are described further below.

**Table 1.** Proportion of SMs associated with TEs across MA lines of two C. reinhardtii strains. Note that one translocation involved both CryptonF-1_cRei and Un3/Un12, and the number of TEs does not always match the number of TE-mediated SMs.

| Strain | SM | % associated with TEs (count/total) | TE families (count) |
| --- | --- | --- | --- |
| CC-1952 | Deletion | 33.33 (1/3) | *Dualen-4b_cRei* (1) |
|  | Duplication | 86.67 (13/15) | *Dualen-4b_cRei* (13) |
|  | Inversion | 0 (0/1) | NA |
|  | Translocation | NA | NA |
| CC-2931 | Deletion | 13.33 (2/15) | *CryptonF-1_cRei* (2) |
|  | Duplication | 2.27 (1/44) | *Dualen-4b_cRei* (1) |
|  | Inversion | 64.28 (9/14) | *CryptonF-1_cRei* (8); *Un3/Un12* (1) |
|  | Translocation | 75.00 (15/20) | *CryptonF-1c_Rei* (8); *Un3/Un12* (8) |

**Table 2.** Proportion of SMs associated with homology-based mechanisms in MA lines of two C. reinhardtii strains.

| Strain | SM | % with macrohomology<br>(count/total) | % with microhomology<br>(count/total) |
| --- | --- | --- | --- |
| CC-1952 | Deletion | 0 (0/3) | 33.33 (1/3) |
|  | Duplication | 6.67 (1/15) | 0 (0/15) |
| CC-2931 | Deletion | 0 (0/15) | 26.67 (4/15) |
|  | Duplication | 4.54 (2/44) | 6.82 (3/44) |

Rearrangements mediated by *Un3/Un12* generally featured breakpoints that coincided with the 3’ end of the transposons, most frequently repaired so that one derived chromosome featured two elements in a tail-to-tail arrangement. For example, this outcome was observed in a reciprocal translocation between chromosomes 14 and 17 in CC-2931 L9 (Fig. 4A). While we do not yet understand the mechanism of these newly discovered TEs, the presence of target site duplications suggest that they may cause DSBs, and their simultaneous insertion at different genomic regions could lead to aberrant repair and rearrangements. *CryptonF-1_cRei*-mediated rearrangements were associated with both insertions and excisions. For example, CC-2931 L11 experienced a reciprocal translocation between chromosomes 9 and 11 involving insertions at each breakpoint (Fig. 4B). Conversely, other events involved only one insertion, although in all cases we observed a “CAYCG” target site at the other breakpoint. Notably, the high insertion rates of both *CryptonF-1_cRei* and *Un3/Un12* in CC-2931 L13 (Fig. 3A) resulted in a highly derived karyotype in this line (Fig. S6A). This included a cluster of non-reciprocal translocations involving four chromosomes, mediated by three *Un3* and one *Un12* insertions that may have occurred simultaneously .

**Figure 4.**
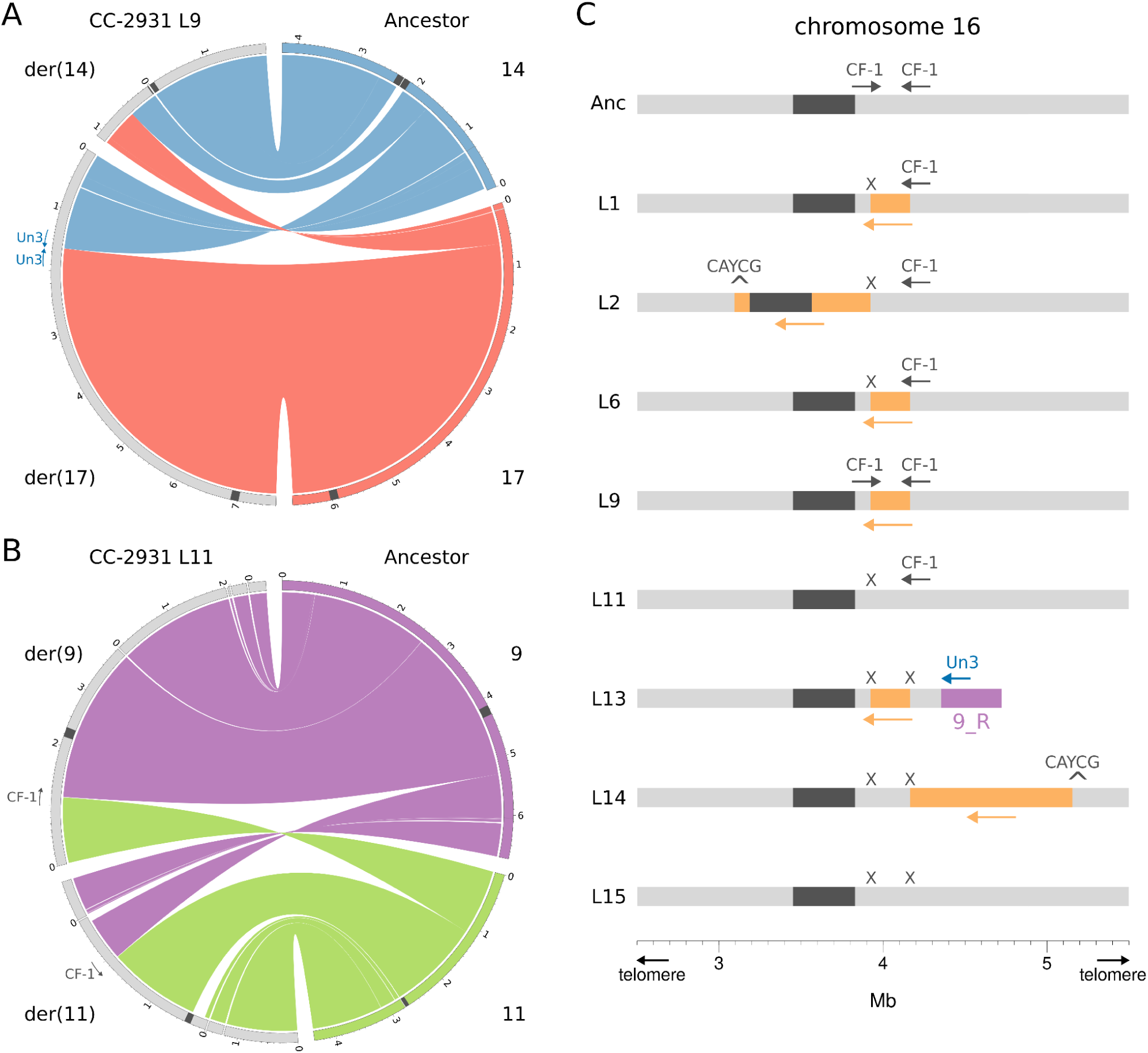
Genomic rearrangements mediated by TEs. Translocations in CC-2931 MA lines 9 (A) and 11 (B). The ancestor chromosomes are represented on the right half of each Circos plot, while the MA line chromosomes (as contigs) are on the left. Chromosome numbers are given for the ancestor, while derived chromosomes (denoted with “der”) are provided for the MA lines based on centromere annotation (dark grey regions). Scale is in megabases. CF-1 indicates *CryptonF-1_cRei* insertions. The direction of the arrows indicates the 5’ to 3’ orientation of the TE sequence. C) *CryptonF-1_cRei* (CF-1) mediated inversions on chromosome 16 in CC-2931 MA lines. Ancestor and MA line genomes are represented from top to bottom. The dark grey region represents the centromere, while the orange blocks represent inversions. The purple region in L13 shows an *Un3*-mediated translocation with chromosome 9. Grey arrows indicate the orientation of *CryptonF-1_cRei* from 5’ to 3’, while ‘X’ indicates *CryptonF-1_cRei* excisions.

Remarkably, *CryptonF-1_cRei* was associated with six inversions in different MA lines on chromosome 16 (Fig. 4C). In the CC-2931 ancestor, chromosome 16 featured two *CryptonF-1_cRei* copies in opposite orientation separated by ∼233 kb. We did not observe this ancestral state in any of the eight MA lines. The ∼233 kb region was inverted in four MA lines, with either none, one or both of the copies excised. In L2 and L14 longer inversions were mediated between one of the *CryptonF-1_cRei* copies and independent “CAYCG” target sites elsewhere on the chromosome (L2 ∼830 kb between copy 1 and an upstream target site; L14 ∼991 kb between copy 2 and a downstream target site). The presence of target sites at each breakpoint (featuring either an insertion or excision, or neither) in both inversions and translocations may suggest a role for the site-specific recombination activity of the Crypton tyrosine recombinase enzyme in mediating rearrangements.

It is also interesting to note that the CC-2931 ancestral genome has a rearranged karyotype relative to CC-1952 and the standard reference genome, featuring two reciprocal translocations (Craig et al. 2021). The breakpoints of these rearrangements were also associated with *Un3* and *Un12* insertions, implying that these elements have been historically active either in the field or in the laboratory since isolation.

Finally, we observed duplications associated with the LINE *Dualen-4b_cRei*, which was most active in CC-1952 (Fig. 3A), but also caused a single insertion (and associated duplication) in CC-2931. More than half of the *Dualen-4b_cRei* insertions were associated with duplications >50 bp in length, representing 87% of duplications in CC-1952 (Table 1). These duplications (of median length 900 bp, Fig. S7) resembled the variable length target site duplications that flank insertions of non-LTR elements (LINEs and PLEs), i.e. the duplicated sequence flanked either side of the *Dualen* insertions. Such target site duplications are caused by resolution of the DNA nicks introduced during insertion, the distance between cleavage sites corresponding to target site duplication length . These putative *Dualen-4b_cRei* target site duplications are considerably longer than other large target site duplications reported previously, e.g. the 126 bp target site duplications observed in *R9* LINEs of rotifers (Gladyshev and Arkhipova, 2009). Active *Dualens* have not been observed previously, and these exceptionally long target site duplications are possibly mediated by the dual action of RLE and APE endonucleases (Fig. 3B), which are uniquely present together in the *Dualen* clade (Kojima and Fujiwara 2005).

### Homology-mediated structural mutations

We next attempted to identify homology-based DSB repair mechanisms that may have mediated the remaining SMs. These included most deletions and duplications in CC-2931 (Table 2). Many mechanisms associated with SMs involve DSB repair, which typically proceeds via two distinct pathways: homologous recombination and canonical non-homologous end joining. As previously mentioned, homologous recombination can induce SMs via recombination between interspersed paralogous sequences (i.e. via non-allelic homologous recombination). Non-homologous end joining can also mediate SMs. For example, two DSBs present on the same chromosome could be repaired aberrantly, yielding a deletion or inversion, or two DSBs present on different chromosomes could yield a translocation. Several, other DSB repair pathways that rely on varying lengths of homology tracts also exist (So et al. 2017). For example, both microhomology-mediated end joining (∼1-16 bp microhomology) and single-strand annealing (>30 bp macrohomology) inherently generate deletions at single DSBs, since they involve a process where DNA ends are degraded to single-strand sequences (i.e. DNA end resection) to reveal homology, followed by deletion of any sequence overhanging the homology tract (i.e. heterologous flaps) (Sfeir & Symington, 2015).

We found evidence of macrohomology (>30 bp) for only 5.1% of duplication events, one event in CC-1952 and two in CC-2931 (Table 2, Fig. S8), which could potentially have been caused by non-allelic homologous recombination. In all three of these duplications, the paralogous sequence did not involve TEs. We found no evidence of macrohomology-mediated deletions, suggesting little role for single-strand annealing in the mediation of SMs. In contrast, we identified microhomologies in 27.8% of deletions, four in CC-2931 and one in CC-1952 (Fig. 4A, Fig S.9). An interesting example of this phenomenon occurred in CC-2931 L6, where we observed several clustered SNMs and INDELs flanking a deletion (Fig. S10), a phenomenon that has been observed flanking DSBs repaired by microhomology-mediated end joining (Sinha et al. 2017). Although such hypermutability was not observed for the other microhomology-associated deletions, they were generally shorter than deletions where we detected no homology tracts (median length 225 bp vs. 2,945 bp; W test, P = 3.77 x 10^-2^). The shorter length of these deletions may be consistent with DNA end resection involved in microhomology-mediated end joining.

We also found three duplications in CC-2931 that showed five, three and two nucleotides of sequence homology between their breakpoints. Although these could be consistent with microhomology-mediated mechanisms, such homology may simply have occurred by chance: when we calculate the expectation of such short homology from the random sampling of 20 bp sequences, the observed frequency of microhomologies is not higher than the random expectation. Finally, we found no evidence of either micro- or macrohomology at the breakpoints of the inversions and translocations not associated with active TEs. However, we observed short deletions at several of the breakpoints of these rearrangements, potentially consistent with repair via non-homologous end joining. Notably, 40% of non-TE associated inversions and translocations occurred in a single CC-2931 line, L15 (Fig. S6B).

### Tandem repeat mutations

Tandemly repeated sequences are known to be hypermutable and evolve via mechanisms that include replication slippage and unequal exchange (Lower et al. 2018). As mentioned above, unlike most other SMs, tandem repeat mutations (TRMs, grouping expansions and contractions >50 bp) occurred at similar rates in CC-2931 and CC-1952 MA lines, perhaps due to the independence of their underlying mechanisms compared to other SMs. Approximately 22% of TRMs occurred in microsatellites (tandem repeats with monomers <10 bp), and most of the remainder occurred in satellite DNA. It is important to note that the rate of TRMs per site per generation is in fact several times higher than the genome-wide rate displayed in Figure 2B, since TRMs by definition can only occur in the tandem repeats (∼12% of the genome), effectively resulting in a smaller ‘callable’ genome.

We also observed TRMs at the sequences that constitute the *C. reinhardtii* centromeres and subtelomeres. The centromeres mostly consist of an *L1* LINE retrotransposon, *ZeppL-1_cRei* (Craig et al. 2021a), and we observed expansions and contractions of these sequences (generating 7.0% of TRMs), consistent with centromere evolution via satellite-like mechanisms (e.g. unequal exchange) rather than active transposition. Subtelomeres feature a unique satellite called *Sultan*, and we observed expansions and contractions of the *Sultan* monomer within the same subtelomere, as hypothesised by Chaux-Jukic et al. (2021). Interestingly, we observed one example in each of CC-1952 (∼94 kb) and CC-2931 (∼163 kb) of the truncation of a chromosome terminus, followed by *de novo* telomere addition (i.e. “telomere healing”, Fig. S12). A small number of TRMs involved the expansion or contraction of tandemly repeated gene families. Examples included 5S ribosomal RNA arrays and clusters of the large *Chlamydomonas*-specific *NCL* gene family on chromosome 15, which encode RNA binding proteins and appear to be experiencing rapid and ongoing evolution in *C. reinhardtii* (Boulouis et al. 2015).

Finally, we observed one example of a “mobile” satellite, which caused four insertions ranging from ∼0.5 to >21 kb in three different CC-2931 MA lines. This satellite, *MSAT-11_cRei*, consists of an ∼1.9 kb monomer and does not feature any characteristics typical of a TE. We have recently observed mobile insertions of *MSAT-11_cRei* in other *C. reinhardtii* strains (CC-1690 and CC-4532, Craig et al. in prep.). Although mechanisms mediating satellite dissemination are not well understood, it has been observed in other species, and may be an important mechanism in satellite evolution (Ruiz-Ruano et al. 2016).

## Discussion

In total, we identified 563 SMs in 12 MA lines derived from the *C. reinhardtii* strains CC-1952 and CC-2931. This provides, to our knowledge, the first direct estimates of the rates and spectra of *de novo* SMs based on long-read sequencing of MA lines. In agreement with previous results on the rates of SNMs and INDELs (Ness et al. 2015), the rates and spectra of SMs vary greatly between MA lines and strains. SMs represented a substantial proportion of the overall genomic mutation rate in both strains (6% in CC-1952, and 12% in CC-2931).

### Calling structural mutations from long-reads and assemblies

*De novo* mutations are inherently rare events, and methods for their detection must therefore be highly accurate to reliably estimate their rates and spectra. Our ability to detect SMs was aided by major advances in both long-read sequencing technology and structural variant callers. Nonetheless, our results show that the detection of structural variants remains much more challenging than the detection of shorter variants, making it necessary to use a combination of approaches. Because there was only partial overlap between variants identified by different callers, it is possible that some SMs were undetected in our study. While a read-based caller (Sniffles) was indispensable for the identification of many variants (especially duplications), this approach failed to detect the full range of SMs, and methods based on genome assembly were also required. The pan-genome approach implemented by Cactus and vg was particularly successful, calling almost 80% of all curated SMs.

However, in all cases, our results demonstrated that structural variant callers are likely to yield high rates of false positives, even when the analysed samples are nearly genetically identical, as in our experiment. Although we curated many genuine SMs in tandem repeats, these regions appeared to be responsible for the majority of false positive calls. Accurately calling SMs in tandem repeats may require specific tool development, and given that manual curation of variants is unlikely to be manageable in larger and more complex genomes than that of *C. reinhardtii*, masking of tandem repeats (including relatively simple microsatellites) may be appropriate in automated analyses. The substantial contribution of assembly errors to rejected calls may also warrant the use of multiple assemblers in variant calling, or the development of methods that combine assembly-based structural variant detection with read-based verification. Overall, we recommend sequencing samples with sufficient coverage to enable *de novo* assembly where possible, followed by both assembly and read-based structural mutation or variant calling. If possible, manual verification with visualisation tools such as IGV should also be performed.

### Mechanisms underlying structural mutation and between strain heterogeneity in rates and spectra

Excluding expansions and contractions of tandem repeats, ∼79% of SMs were associated with TEs. Of these SMs, 84% were TE insertions and excisions, which formed major components of the SM spectra in both strains. The remaining proportion involved other types of SMs putatively mediated by active TEs, including the majority of inversions and translocations in CC-2931, and almost all duplications in CC-1952. The *C. reinhardtii* genome contains more than 200 TE families from a diverse array of subclasses and superfamilies, almost all of which show evidence of recent activity (Craig 2021). However, none of the three TEs implicated in mediating rearrangements or duplications have previously been observed as active elements: Cryptons (Goodwin et al. 2003) and *Dualen* LINEs (Kojima and Fujiwara 2005) were first described from multi-copy repeats in genetic and genomic data, while the Rep-encoding *Un12* element found here has not been described previously.

Beyond TEs, we generally found little role for homology-based mechanisms of DSB repair, with the possible exception of the microhomology-mediated end joining pathway in mediating deletions of moderate lengths. This is in stark contrast to studies of SM in yeast, where non-allelic homologous recombination has been shown to be the predominant mechanism mediating deletions, duplications and rearrangements (Sui et al. 2020). Similarly, approximately 10-20% of *de novo* SMs in humans are thought to be mediated by non-allelic homologous recombination (Parks et al. 2015). Instead of homology-based mechanisms, the non-homologous end joining repair pathway may have been involved in many of the SMs not mediated by TEs. This result is consistent with the very low rates of homologous recombination observed in *C. reinhardtii* under vegetative growth (where the species is haploid), where non-homologous end joining is the dominant DSB repair pathway (Ferenczi et al. 2021). The relative rates of non-homologous end joining and homologous recombination in the repair of DSBs have been implicated in many aspects of genome evolution, such as the evolution of base composition (Weissman et al. 2019) and intron density (Farlow et al. 2011), and it is likely that variation in the activity of different DSB repair pathways also leads to substantial differences in SM spectra among species.

In our previous analysis (Ness et al. 2015) and re-analysis herein, which employed Illumina and PacBio HiFi reads, respectively, we found substantial variation in the rates of SNMs and INDELs between the CC-1952 and CC-2931 strains. Nucleotide diversity among *C. reinhardtii* strains is high (i.e. π = ∼2-3%; Flowers et al. 2015, Craig et al. 2019), and some of the within-species variation in mutation rates may be caused by presence of mutator alleles in certain strains. Consistent with SNM and short INDEL mutations, the SM rate in CC-1952 MA lines was significantly lower than in CC-2931 MA lines. Furthermore, we here found that the SM spectra also differed substantially between the strains. CC-2931 has a higher overall rate of transposition than CC-1952, involving a more diverse array of TEs. As a result, CC-2931 MA lines experienced many TE-mediated inversions and translocations that were absent in CC-1952. TE suppression is not fully understood in *C. reinhardtii*, but likely occurs at transcriptional and post-transcriptional levels (van Dijk et al. 2006) via mechanisms including repressive histone modifications (Jeong et al. 2002; Zhang et al. 2002) and RNA interference (Casas-Mollano et al. 2008). Since the CC-2931 and CC-1952 genomes both harbour many more potentially active TE families than we observed (see above), all but a few families appear to be silenced effectively. TE suppression could conceivably differ between CC-2931 and CC-1952 due to genetic variation in genes involved in silencing pathways. Alternatively, environmental factors could impact TE activity. Almost nothing is known about local adaptation in *C. reinhardtii*, although natural isolates differ in their growth rates and fitness estimates under laboratory conditions (Morgan et al. 2014; Kraemer et al. 2017). CC-1952 and CC-2931 were sampled almost 1,600 kilometres apart (from Minnesota and North Carolina, respectively), and it is possible that they differ in their extent of adaptation to the highly artificial laboratory environment. Such variation could potentially cause stress-related interactions with transposition, which can be complex, and result in both TE activation or repression (Horváth et al. 2017). Incidentally, although nothing is known about their rates of transposition, a similar number of TE families to that which we observed in CC-2931 are known to be active in *C. reinhardtii* ‘laboratory strains’ (a collection of related strains that almost all *C. reinhardtii* research is performed on), although all of the TEs active in laboratory strains (e.g. *Gulliver, TOC1, MRC1, Tcr1* and *Tcr3*; Craig 2021) differ from those that are active in CC-2931. Therefore, CC-1952 may be exceptional in having few active TE families, compared to other *C. reinhardtii* strains.

Appart from a higher rate of transposition, CC-2931 MA lines also may have experienced a higher rate of DSBs than CC-1952 MA lines, which may explain the higher rates of duplications, deletions and rearrangements, even after TE-mediated SMs have been accounted for. DSBs are generally considered the most mutagenic DNA lesions (So et al. 2017) and are induced by intrinsic cellular factors (e.g. replicative and oxidative stresses) and by exogenous sources (e.g. mutagens). The rate of DSBs in *C. reinhardtii* is, however, not well-understood, and differences between the strains could arise from genetic or environmental factors.

### Evolutionary implications of high and variable rates of structural mutation

Population-level long-read sequencing projects have generally found that structural variants are prevalent, yet segregate at low frequencies, implying that most are strongly deleterious (Chakraborty et al. 2019; Weissensteiner et al. 2020). Although we have not explored the relationship between SMs and fitness, the genomic distribution of SMs suggests that many will have large fitness effects. We found that large deletions and duplications affected coding and other exonic sequences at similar rates to their genomic distribution. In contrast, a growing body of evidence suggests that coding sequences are less mutable than intergenic sequences with respect to SNMs and INDELs (Lee et al. 2012; Belfield et al. 2018; Krasovec et al. 2017; López-Cortegano et al. 2021; Monroe et al. 2022), presumably due to improved efficiency of the DNA repair machinery or the presence of epigenetic factors affecting coding sequences. Furthermore, although certain TE families exhibited insertion biases for introns (*Chlamys* PLEs) or intergenic sequence far from genes (*EnSpm* DD(E/D) transposons), overall TEs exhibited a bias towards 5’ UTRs and gene proximal intergenic regions, as has been observed for various other TEs (Zhang et al. 2020). Gene proximal TE insertions can have important effects on gene expression, for example via the disruption of regulatory sequences, regional effects of transcriptional silencing, or the deposition of new regulatory elements by the TE (Cridland et al. 2015; Uzunović et al. 2019; Rech et al. 2022). The breakpoints of chromosomal rearrangements (inversions and translocations) also had a similar genomic distribution to that of TEs, due to their association. Although not a factor in our experiment, translocations may also yield chromosomal imbalances and have substantial deleterious effects in meiosis. Taken together with the relatively high fraction of the total mutation rate explained by SMs herein, the genomic distribution of SMs implies that SMs may contribute a substantial mutation load and potential for adaptive evolution.

Our results particularly highlight the prevalence and importance of TEs among SMs, and support a prominent evolutionary role for TEs. As discussed, the heterogeneity in the rate and identity of TE insertions between CC-2931 and CC-1952 contributed substantially to the overall differences in SM rates and spectra between strains. Our results also suggest that different species, populations and even individuals may differ considerably in their SM spectra as a result of their active repertoire of TEs. More work is required to investigate the rate of new SMs in other species, elucidating the phylogenetic generality of results observed here. In particular, it will be important to test whether similar within species variation in SM rates and spectra exists in other taxa. We hope that our results add to the weight of evidence supporting the importance and prevalence of SMs, and further encourage the ongoing movement towards structural variant discovery via assembly-based methods.

## Methods

### Biological samples and nucleic acids extraction

The MA line ancestors were *Chlamydomonas reinhardtii* wild strains CC-1952 (from Minnesota, 1986) and CC-2931 (North Carolina, 1991), which were originally obtained from the Chlamydomonas Resource Center (https://www.chlamycollection.org/). The MA experiment was conducted by Morgan et al. (2014). Briefly, MA lines were initiated from the ancestor strains, and cultured on Bold’s medium agar plates under white light at 25℃. MA lines were bottlenecked at regular intervals of 3 to 5 days by randomly picking single colonies and transferring them from one plate to another, and maintaining them for ∼1,000 generations (within-strain mean numbers of generations were 1,066 for CC-1952 and 1,050 for CC-2931), after which Illumina sequencing was performed. The original ancestors and MA lines from the end of the experiment were cryopreserved in liquid nitrogen.

For this study, we revived the CC-1952 and CC-2931 ancestors along with several MA lines from cryopreservation, and grew all samples in liquid Bold’s medium before transferring to agar slants in order to produce stock cultures. Four CC-1952 MA lines (L1, L3, L6 and L15) and eight CC-2931 MA lines (L1, L2, L6, L9, L11, L13, L14 and L15) were selected for sequencing together with the two ancestors. Cells were inoculated in 6-well plate liquid cultures and grown for 4 days under constant light to produce sufficient biomass for DNA extraction. High molecular weight genomic DNA was extracted using a cetyltrimethylammonium bromide and phenol:chloroform protocol, following Craig et al. (2021a).

RNA was extracted in triplicate from independent cultures of the CC-2931 ancestor grown in liquid Bold’s medium under constant light via a Maxwell RSC 48 instrument.

### Nucleic acids sequencing

All of the CC-1952 samples and six of eight CC-2931 MA lines were sequenced on the PacBio Sequel II platform with a 30hr movie, using the circular consensus sequencing (CCS) mode to generate HiFi reads. Samples were multiplexed, with between four and six samples per SMRT cell. Library and barcoding preparation, sequencing, CCS analysis and demultiplexing were performed at the Earlham Institute (Norwich, United Kingdom). Mean read length was ∼20 kb and mean coverage ∼30x per sample.

The CC-2931 ancestor and CC-2931 MA lines L2 and L9 were sequenced on individual SMRT cells using the PacBio Sequel I platform with a 10hr movie, using the continuous long read (CLR) mode. Library preparation and sequencing was performed at Edinburgh Genomics (Edinburgh, United Kingdom). Mean read length was ∼12 kb and mean coverage was ∼55x per sample.

CC-2931 ancestor RNA-seq library preparation was conducted with the NEB mRNA stranded library preparation kit and Illumina NovaSeq preparation. The three replicate samples were sequenced using Illumina 100 bp paired-end sequencing. Library preparation and sequencing were performed by Genome Quebec.

### Genome assembly

MA lines were assembled *de novo* to the contig-level. Flye v2.8.2 (Kolmogorov et al. 2019) was selected for assembly, since it produced assemblies that were most representative of the haploid state (i.e. other assemblers yielded redundant contigs in repetitive regions). A genome length of 111.1 Mb was assumed (-g 111.1m in Flye). Post-processing was performed with purge_dups (Guan et al. 2020). Error correction was only performed for the two CC-2931 MA lines (L2 and L9) sequenced using CLR, by applying two iterative rounds of the Arrow algorithm (Hepler et al. 2016). The MA lines ancestors were assembled *de novo* to the chromosome-level using a combination of assembly methods (see Supplemental Material).

Assembly metrics were quantified using QUAST v5.0.2 (Gurevich et al. 2013). Assembly completeness was estimated using BUSCO v4.0.6 (Manni et al. 2021), which was run in genome mode using the chlorophyta_odb10 dataset (“augustus_species chlamy2011”).

### Genome annotation

Structural annotation of the CC-2931 ancestor assembly was performed using BRAKER v2.15 (Brůna et al. 2021). RNAseq data were mapped to the assembly using STAR v2.7.9 (“--alignIntronMax 5000 --twopassMode Basic”) (Dobin et al. 2013). BRAKER2 was first run on an assembly softmasked for all repeats (see below) using existing Augustus parameters for *C. reinhardtii* (“--species=chlamy2011 --skipAllTraining”). A second BRAKER2 run was performed with UTR prediction (“--stranded=+,- --UTR=on”), where the input BAM alignments were split to forward and reverse strand read sets using samtools v1.9 (Danecek et al. 2021). The run without UTR prediction was used as the primary annotation based on a superior BUSCO score (protein mode). However, for gene models where coding sequence coordinates had a one-to-one correspondence between the two runs (with variation permitted at only one exon for models with >2 exons) the model with UTRs was introduced as a replacement. Gene models were filtered if their coding sequence had a >30% intersect with transposable element (TE) sequence or a >70% intersect with simple repeats (i.e. microsatellites) identified with RepeatMasker v4.0.9 (http://www.repeatmasker.org).

Repeat annotation was performed for the CC-2931 and CC-1952 ancestor assemblies. A custom library of *C. reinhardtii* TEs and satellites (Craig 2021) was first passed to RepeatMasker. A small number of TE families newly identified in this study were manually curated and added to the library using methods described by Goubert et al. (2022). The RepeatMasker annotations were supplemented with additional microsatellites and satellites identified by Tandem Repeats Finder (“2 7 7 80 10 50 1000 -f -d -m -ngs”) (Benson 1999). A final set of tandem repeats for each assembly was produced by combining simple repeats and satellites from RepeatMasker, satellites and microsatellites from Tandem Repeats Finder, and manually curated centromeric, subtelomeric and ribosomal DNA array coordinates. Centromeres were identified based on the span of the constituent LINE *ZeppL-1_cRei* (Craig et al. 2021a) and subtelomeres as any sequence telomere-proximal to the characteristic *spacer* sequence (see Chaux-Jukic et al. 2021). A microsatellite was considered as a tandem repeat sequence with a monomer length <10 bp, with tandem repeats consisting of any longer monomer length considered satellite DNA.

### Mapping and alignment

MA line PacBio read mapping was performed against the appropriate reference assembly using pbmm2 (https://github.com/PacificBiosciences/pbmm2), adjusting the “--preset” flag for HiFi when appropriate. To avoid false positives when calling variants, reads with tags indicating secondary alignments were removed using samtools (Danecek et al. 2021).

MA line assemblies were aligned to the appropriate reference assembly using minimap2 (“-x asm5”) (Li 2018). For visualisation of certain tandem repeat mutations, unimap v0.1-r41 (https://github.com/lh3/unimap), a tool derived from minimap2 and optimised for assembly alignments, was also used.

A pangenome alignment was produced for each ancestor and its MA lines using Cactus v1.3.0 (Armstrong et al. 2020). All assemblies were first softmasked for repeats that had been first identified by RepeatMasker and Tandem Repeats Finder (see above). Minigraph v0.15-r426 (“-xggs”) (Li et al. 2020) was used to produce the GFA (Graphical Fragment Assembly) file that was provided as input for cactus-graphmap together with softmasked assemblies. The resulting PAF file was then passed to cactus-align (“--pangenome –pafInput --outVG”) in order to produce the final pangenome in VG (variation graph) format.

### Callable sites

Callable sites were defined as genomic sites where structural mutations (SMs) could be called with high confidence based on visualisation of the sequencing and assembly data. We previously defined callable sites based on short-read mapping parameters (Ness et al. 2015; López-Cortegano et al. 2021). Here, we used criteria based on the *de novo* genome assemblies of the ancestors and MA lines. First, the ancestor assembly was aligned against itself with minimap2 (“-x asm5”), and genomic regions that were unrepresented in the PAF file (i.e. that were mismapped) were deemed as uncallable (typically the most repetitive regions). Then, a similar procedure was followed for each MA line, by aligning their assemblies to the ancestor genome, extracting and merging mapping coordinates from the PAF file, and identifying unmapped regions. These regions were intersected using bedtools v2.30.0 (Quinlan and Hall, 2010), so that unmapped regions present in more than one line could be identified as uncallable (since lack of mapping in a single line could be an SM, such as a large deletion). Finally, a few such regions were manually reincluded as callable, based on visualisation of assembly mappings in IGV (e.g. a region containing a DNA transposon that was excised, and hence did not map, in multiple lines).

To compare callable sites between our previous Illumina sequencing and the PacBio sequencing described here, callable site coordinates from Ness et al. (2015) were converted to the new ancestor assemblies. A whole-genome alignment of the v5 reference assembly and the ancestor assemblies was generated using Cactus (Armstrong et al. 2020). Coordinates were then lifted over to the relevant ancestor assembly using the HAL tools command halLiftover (Hickey et al. 2013).

### Structural mutation identification

SMs were called using three independent pipelines. Sniffles v1.0.12b (Sedlazeck et al. 2018) was used to call SMs based on read alignments. BAM files were preprocessed using samtools-calmd to generate the MD tag, which provides information on mismatching positions (i.e. variable coordinates in the reads). Sniffles was first run on each MA line individually, and the resulting VCF files were merged using SURVIVOR v1.0.7 (Jeffares et al. 2017). Sniffles was then run again for all MA lines using the option “--Ivcf”, using the merged VCF as input and enabling the calling of SMs that could be present in more than one MA line. SURVIVOR was then used again to generate a multi-sample VCF.

MUM&Co v3 (O’Donnell and Fischer 2020) was used to call SMs from individual alignments of MA line assemblies to their ancestral reference. MUM&Co calls variants based on alignments produced by MUMmer v4 (Marçais et al. 2018). A genome size of 110 Mb was used (“-g 110000000”) and variants were obtained as TSV and VCF files. vg (variation graph tool; Garrison et al. 2018) was used to call variants directly from the pangenome alignments using the deconstruct command. The resulting VCF file for each strain was reduced to variants >50 bp.

All called variants in callable regions were manually curated via visualisation of read and assembly alignments using IGV (Robinson et al. 2011). SMs were rejected if they were not supported unambiguously by the read alignments. Complex SMs, including large rearrangements and duplications, were further visualised using Ribbon v1.1 (Nattestad et al. 2021). Duplications and deletions were curated as tandem repeat expansions or contractions if they involved the duplication or deletion of one or more monomers of a tandem repeat. Most fell within existing tandem repeat annotations (satellites and microsatellites, see above), while a small number required manual inspection of insertion/deletion flanks by self-vs-self dotplots generated using the MAFFT v7 online server (Katoh et al. 2019). Deletions that perfectly intersected with TEs annotated by RepeatMasker were called as mobile excisions. Mobile insertions were identified by using the inserted sequence as a blastn (Camacho et al. 2009) query against the repeat library and manually inspecting the output. See Figures S11-S25 for examples of SM visualisation.

Variants >30 kb (approximately the upper limit of read lengths), including large inversions and translocations, were manually curated based on the MA line assembly minimap2 PAF alignments files (Supplementary S7). This process led to the identification of a small number of additional large SMs that were not called by any of the three pipelines. Each inversion and translocation breakpoint was manually checked for the presence of TE insertions, which could also be visualised with Ribbon. Five rearrangement SMs could not be fully characterised (e.g. if one breakpoint was clearly supported but the second was in an uncallable region) and were arbitrarily included as translocations.

### SNMs and INDELs

Deepvariant 1.1.0 (Poplin et al. 2018) was used for calling SNMs and INDELs based on read alignments, setting the option “--model_type=PACBIO”. Deepvariant was run on individual MA line alignment files. The resulting VCF files were merged using GLNexus 1.3.1 (“ --config Deepvariant_unfiltered”) (Yun et al. 2020). The merged VCF file was further processed to retain only high quality calls (QUAL ≥ 20) of sites called as homozygous genotype (deepvariant assumes diploid genomes) and biallelic variants, with a minimum read depth of 8. In addition, only variant calls that were unique to a single MA line were retained as mutations. All SNMs and INDELs were confirmed visually from IGV snapshots.

### Genomic distributions of SMs and TE insertions

The coordinates of SMs in CC-2931 were intersected with genomic annotations (coding sequence, introns, etc.) using bedtools. Tandem repeat mutations (TRMs) and excisions were not analysed. Additionally, we also calculated the extent of overlap of deletions, duplications and inversions based on the entire span of these SMs. The observed distributions of SMs were compared against null expectations based on random sampling of the callable genome. For the analysis of individual TE families the random expectation was adjusted to account for insertion target sequences (see main text). The target motifs were identified using SeqKit v2.1.0 (Shen et al. 2016) and these motifs were then sampled from the callable genome.

### Sequence homology

To explore the role of different DSB repair pathways in SM generation, we searched for patterns of micro and macrohomology at SM breakpoints, following Belyeu et al. (2021). We first searched for evidence of macrohomology between the breakpoints of deletions, duplications, inversions and translocations that were not previously associated with TEs (≥95% sequence identity detected by the megablast; Camacho et al. 2009). Query and target sequences were the 100 bp upstream and downstream from each breakpoint. In addition to macrohomology, we looked for patterns of microhomology. Given that microhomology mechanisms such as microhomology-mediated end joining can require as little as 2 bp of homology, we manually investigated the 20 bp of sequence surrounding breakpoints.

## Acknowledgements

We thank Alexander Suh and Aaron Vogan for valuable discussions on TE classification and mechanisms. Next-generation sequencing and library construction was delivered via the BBSRC National Capability in Genomics and Single Cell Analysis (BBS/E/T/000PR9816) at the Earlham Institute by members of the Genomics Pipelines Group. Analyses performed here made use of the high-performance computing resources at the Edinburgh Compute and Data Facility. This project has received funding from the European Research Council under the European Union’s Horizon 2020 research and innovation programme (grant agreement no. 694212).

## Data availability

Raw reads and genome assemblies will be deposited at the NCBI Sequence Read Archive (SRA) under BioProject PRJNA839925.

## Notes

### Competing Interest Statement

The authors have declared no competing interest.

## References

1. Armstrong J, Hickey G, Diekhans M, Fiddes IT, Novak AM, Deran A, Fang Q, Xie D, Feng S, Stiller J et al. 2020. Progressive Cactus is a multiple-genome aligner for the thousand-genome era. Nature 587: 246–251.

2. Bateman AJ. 1959. The viability of near-normal irradiated chromosomes. Int J Radiat Biol 1: 170–180.

3. Belfield EJ, Ding ZJ, Jamieson FJC, Visscher AM, Zheng SJ, Mithani A, Harberd NP. 2018. DNA mismatch repair preferentially protects genes from mutation. Genome Res 28: 66–74.

4. Belyeu JR, Brand H, Wang H, Zhao X, Pedersen BS, Feusier J, Gupta M, Nicholas TJ, Brown J, Baird L et al. 2021. De novo structural mutation rates and gamete-of-origin biases revealed through genome sequencing of 2,396 families. Am J Hum Genet 108: 597–607.

5. Benson G. 1999. Tandem repeats finder: a program to analyze DNA sequences. Nucleic Acids Res 27: 573–580.

6. Böndel KB, Kraemer SA, Samuels T, McClean D, Lachapelle J, Ness RW, Colegrave N, Keightley PD. 2019. Inferring the distribution of fitness effects of spontaneous mutations in *Chlamydomonas reinhardtii*. PLoS Biol 17: e3000192.

7. Böndel KB, Samuels T, Craig RJ, Ness RW, Colegrave N, Keightley PD. 2021. The distribution of fitness effects of spontaneous mutations in *Chlamydomonas reinhardtii* inferred using frequency changes under experimental evolution. Biorxiv doi:https://doi.org/10.1101/2021.09.29.462298.

8. Boulouis A, Drapier D, Razafimanantsoa H, Wostrikoff K, Tourasse NJ, Pascal K, Girard-Bascou J, Vallon O, Wollman FA, Choquet Y. 2015. Spontaneous dominant mutations in *Chlamydomonas* highlight ongoing evolution by gene diversification. Plant Cell 27: 984–1001.

9. Brůna T, Hoff KJ, Lomsadze A, Stanke M, Borodovsky M. 2021. BRAKER2: automatic eukaryotic genome annotation with GeneMark-EP+ and AUGUSTUS supported by a protein database. NAR Genom Bioinform 3: lqaa108.

10. Camacho C, Coulouris G, Avagyan V, Ma N, Papadopoulos J, Bealer K, Madden TL. 2009. BLAST+: architecture and applications. BMC Bioinformatics 10: 421.

11. Casas-Mollano JA, Rohr J, Kim EJ, Balassa E, van Dijk K, Cerutti H. 2008. Diversification of the core RNA interference machinery in *Chlamydomonas reinhardtii* and the role of DCL1 in transposon silencing. Genetics 179: 69–81.

12. Chakraborty M, Emerson JJ, Macdonald SJ, Long AD. 2019. Structural variants exhibit widespread allelic heterogeneity and shape variation in complex traits. Nat Commun 10: 4872.

13. Chaux-Jukic F, O’Donnell S, Craig RJ, Eberhard S, Vallon O, Xu Z. 2021. Architecture and evolution of subtelomeres in the unicellular green alga *Chlamydomonas reinhardtii*. Nucleic Acids Res doi:10.1093/nar/gkab534.

14. Chuong EB, Elde NC, Feschotte C. 2017. Regulatory activities of transposable elements: from conflicts to benefits. Nat Rev Genet 18: 71–86.

15. Craig RJ. 2021. The evolutionary genomics of *Chlamydomonas*. PhD thesis. University of Edinburgh. http://dx.doi.org/10.7488/era/1603

16. Craig RJ, Böndel KB, Arakawa K, Nakada T, Ito T, Bell G, Colegrave N, Keightley PD, Ness RW. 2019. Patterns of population structure and complex haplotype sharing among field isolates of the green alga *Chlamydomonas reinhardtii*. Mol Ecol 28: 3977–3993.

17. Craig RJ, Hasan AR, Ness RW, Keightley PD. 2021a. Comparative genomics of *Chlamydomonas*. Plant Cell 33: 1016–1041.

18. Craig RJ, Yushenova IA, Rodriguez F, Arkhipova IR. 2021b. An ancient clade of *Penelope*-like retroelements with permuted domains is present in the green lineage and protists, and dominates many invertebrate genomes. Mol Biol Evol 38: 5005–5020.

19. Cridland JM, Thornton KR, Long AD. 2015. Gene expression variation in *Drosophila melanogaster* due to rare transposable element insertion alleles of large effect. Genetics 199: 85–93.

20. Danecek P, Bonfield JK, Liddle J, Marshall J, Ohan V, Pollard MO, Whitwham A, Keane T, McCarthy SA, Davies RM et al. 2021. Twelve years of SAMtools and BCFtools. Gigascience 10.

21. De Coster W, Weissensteiner MH, Sedlazeck FJ. 2021. Towards population-scale long-read sequencing. Nat Rev Genet 22: 572–587.

22. Dobin A, Davis CA, Schlesinger F, Drenkow J, Zaleski C, Jha S, Batut P, Chaisson M, Gingeras TR. 2013. STAR: ultrafast universal RNA-seq aligner. Bioinformatics 29: 15–21.

23. Dobzhansky T, Epling C. 1948. The suppression of crossing over in inversion heterozygotes of *Drosophila pseudoobscura*. Proc Natl Acad Sci U S A 34: 137–141.

24. Faria R, Navarro A. 2010. Chromosomal speciation revisited: rearranging theory with pieces of evidence. Trends Ecol Evol 25: 660–669.

25. Farlow A, Meduri E, Schlotterer C. 2011. DNA double-strand break repair and the evolution of intron density. Trends Genet 27: 1–6.

26. Ferenczi A, Chew YP, Kroll E, von Koppenfels C, Hudson A, Molnar A. 2021. Mechanistic and genetic basis of single-strand templated repair at Cas12a-induced DNA breaks in *Chlamydomonas reinhardtii*. Nat Commun 12: 6751.

27. Feschotte C, Pritham EJ. 2007. DNA transposons and the evolution of eukaryotic genomes. Annu Rev Genet 41: 331–368.

28. Flowers JM, Hazzouri KM, Pham GM, Rosas U, Bahmani T, Khraiwesh B, Nelson DR, Jijakli K, Abdrabu R, Harris EH et al. 2015. Whole-genome resequencing reveals extensive natural variation in the model green alga *Chlamydomonas reinhardtii*. Plant Cell 27: 2353–2369.

29. Garrison E, Siren J, Novak AM, Hickey G, Eizenga JM, Dawson ET, Jones W, Garg S, Markello C, Lin MF et al. 2018. Variation graph toolkit improves read mapping by representing genetic variation in the reference. Nat Biotechnol 36: 875–879.

30. Gladyshev EA, Arkhipova IR. 2009. Rotifer rDNA-specific R9 retrotransposable elements generate an exceptionally long target site duplication upon insertion. Gene 448: 145–150.

31. Goodwin TJ, Butler MI, Poulter RT. 2003. Cryptons: a group of tyrosine-recombinase-encoding DNA transposons from pathogenic fungi. Microbiology 149: 3099–3109.

32. Goubert C, Craig RJ, Bilat A, Peona V, Vogan AA, Protasio AV. 2022. A beginner’s guide to the manual curation of transposable elements Mob DNA 13: 7.

33. Gray YH. 2000. It takes two transposons to tango: transposable-element-mediated chromosomal rearrangements. Trends Genet 16: 461–468.

34. Gregory TR. 2005. Synergy between sequence and size in large-scale genomics. Nat Rev Genet 6: 699–708.

35. Guan D, McCarthy SA, Wood J, Howe K, Wang Y, Durbin R. 2020. Identifying and removing haplotypic duplication in primary genome assemblies. Bioinformatics 36: 2896–2898.

36. Gurevich A, Saveliev V, Vyahhi N, Tesler G. 2013. QUAST: quality assessment tool for genome assemblies. Bioinformatics 29: 1072–1075.

37. Halligan DL, Keightley PD. 2009. Spontaneous mutation accumulation studies in evolutionary genetics. Annu Rev Ecol Evol S 40: 151–172.

38. Han Y, Qin S, Wessler SR. 2013. Comparison of class 2 transposable elements at superfamily resolution reveals conserved and distinct features in cereal grass genomes. Bmc Genomics 14: 71.

39. Hepler NL, Delaney N, Brown M, Smith ML, Katzenstein D, Paxinos EE, Alexander D. 2016. An improved circular consensus algorithm with an application to detect HIV-1 drug resistance associated mutations (DRAMs). Conference on Advances in Genome Biology and Technology.

40. Hickey G, Paten B, Earl D, Zerbino D, Haussler D. 2013. HAL: a hierarchical format for storing and analyzing multiple genome alignments. Bioinformatics 29: 1341–1342.

41. Horváth V, Merenciano M, Gonzalez J. 2017. Revisiting the relationship between transposable elements and the eukaryotic stress response. Trends Genet 33: 832–841.

42. Inaki K, Liu ET. 2012. Structural mutations in cancer: mechanistic and functional insights. Trends Genet 28: 550–559.

43. Jain M, Koren S, Miga KH, Quick J, Rand AC, Sasani TA, Tyson JR, Beggs AD, Dilthey AT, Fiddes IT et al. 2018. Nanopore sequencing and assembly of a human genome with ultra-long reads. Nat Biotechnol 36: 338–345.

44. Jayakodi M, Padmarasu S, Haberer G, Bonthala VS, Gundlach H, Monat C, Lux T, Kamal N, Lang D, Himmelbach A et al. 2020. The barley pan-genome reveals the hidden legacy of mutation breeding. Nature 588: 284–289.

45. Jeffares DC, Jolly C, Hoti M, Speed D, Shaw L, Rallis C, Balloux F, Dessimoz C, Bahler J, Sedlazeck FJ. 2017. Transient structural variations have strong effects on quantitative traits and reproductive isolation in fission yeast. Nat Commun 8: 14061.

46. Jeong BR, Wu-Scharf D, Zhang C, Cerutti H. 2002. Suppressors of transcriptional transgenic silencing in *Chlamydomonas* are sensitive to DNA-damaging agents and reactivate transposable elements. Proc Natl Acad Sci U S A 99: 1076–1081.

47. Joron M, Frezal L, Jones RT, Chamberlain NL, Lee SF, Haag CR, Whibley A, Becuwe M, Baxter SW, Ferguson L et al. 2011. Chromosomal rearrangements maintain a polymorphic supergene controlling butterfly mimicry. Nature 477: 203–206.

48. Kapusta A, Suh A, Feschotte C. 2017. Dynamics of genome size evolution in birds and mammals. Proc Natl Acad Sci U S A 114: E1460–E1469.

49. Katoh K, Rozewicki J, Yamada KD. 2019. MAFFT online service: multiple sequence alignment, interactive sequence choice and visualization. Brief Bioinform 20: 1160–1166.

50. Kazlauskas D, Varsani A, Koonin EV, Krupovic M. 2019. Multiple origins of prokaryotic and eukaryotic single-stranded DNA viruses from bacterial and archaeal plasmids. Nat Commun 10: 3425.

51. Kirkpatrick M. 2010. How and why chromosome inversions evolve. PLoS Biol 8.

52. Kirkpatrick M, Barton N. 2006. Chromosome inversions, local adaptation and speciation. Genetics 173: 419–434.

53. Kojima KK, Fujiwara H. 2005. An extraordinary retrotransposon family encoding dual endonucleases. Genome Res 15: 1106–1117.

54. Kolmogorov M, Yuan J, Lin Y, Pevzner PA. 2019. Assembly of long, error-prone reads using repeat graphs. Nat Biotechnol 37: 540–546.

55. Konkel MK, Batzer MA. 2010. A mobile threat to genome stability: The impact of non-LTR retrotransposons upon the human genome. Semin Cancer Biol 20: 211–221.

56. Krasovec M, Eyre-Walker A, Sanchez-Ferandin S, Piganeau G. 2017. Spontaneous mutation rate in the smallest photosynthetic eukaryotes. Mol Biol Evol 34: 1770–1779.

57. Kraemer SA, Bondel KB, Ness RW, Keightley PD, Colegrave N. 2017. Fitness change in relation to mutation number in spontaneous mutation accumulation lines of *Chlamydomonas reinhardtii*. Evolution 71: 2918–2929.

58. Küpper C, Stocks M, Risse JE, Dos Remedios N, Farrell LL, McRae SB, Morgan TC, Karlionova N, Pinchuk P, Verkuil YI et al. 2016. A supergene determines highly divergent male reproductive morphs in the ruff. Nat Genet 48: 79–83.

59. Kuzmin E, Taylor JS, Boone C. 2022. Retention of duplicated genes in evolution. Trends Genet 38: 59–72.

60. Lee H, Popodi E, Tang H, Foster PL. 2012. Rate and molecular spectrum of spontaneous mutations in the bacterium *Escherichia coli* as determined by whole-genome sequencing. Proc Natl Acad Sci U S A 109: E2774–2783.

61. Li H. 2018. Minimap2: pairwise alignment for nucleotide sequences. Bioinformatics 34: 3094–3100.

62. Li H, Feng X, Chu C. 2020. The design and construction of reference pangenome graphs with minigraph. Genome Biol 21: 265.

63. López-Cortegano E, Craig RJ, Chebib J, Samuels T, Morgan AD, Kraemer SA, Bondel KB, Ness RW, Colegrave N, Keightley PD. 2021. De novo mutation rate variation and its determinants in *Chlamydomonas*. Mol Biol Evol doi:10.1093/molbev/msab140.

64. Lower SS, McGurk MP, Clark AG, Barbash DA. 2018. Satellite DNA evolution: old ideas, new approaches. Curr Opin Genet Dev 49: 70–78.

65. Mahmoud M, Gobet N, Cruz-Davalos DI, Mounier N, Dessimoz C, Sedlazeck FJ. 2019. Structural variant calling: the long and the short of it. Genome Biol 20: 246.

66. Manni M, Berkeley MR, Seppey M, Simão FA, Zdobnov EM. 2021. BUSCO update: novel and streamlined workflows along with broader and deeper phylogenetic coverage for scoring of eukaryotic, prokaryotic, and viral genomes. Mol Biol Evol doi:10.1093/molbev/msab199.

67. Marçais G, Delcher AL, Phillippy AM, Coston R, Salzberg SL, Zimin A. 2018. MUMmer4: A fast and versatile genome alignment system. PLoS Comput Biol 14: e1005944.

68. McClintock B. 1950. The origin and behavior of mutable loci in maize. Proc Natl Acad Sci U S A 36: 344–355.

69. Miga KH, Koren S, Rhie A, Vollger MR, Gershman A, Bzikadze A, Brooks S, Howe E, Porubsky D, Logsdon GA et al. 2020. Telomere-to-telomere assembly of a complete human X chromosome. Nature 585: 79–84.

70. Monroe JG, Srikant T, Carbonell-Bejerano P, Becker C, Lensink M, Exposito-Alonso M, Klein M, Hildebrandt J, Neumann M, Kliebenstein D et al. 2022. Mutation bias reflects natural selection in *Arabidopsis thaliana*. Nature 602: 101–105.

71. Morgan AD, Ness RW, Keightley PD, Colegrave N. 2014. Spontaneous mutation accumulation in multiple strains of the green alga, *Chlamydomonas reinhardtii*. Evolution 68: 2589–2602.

72. Mukai T. 1964. The genetic structure of natural populations of *Drosophila melanogaster*. I. spontaneous mutation rate of polygenes controlling viability. Genetics 50: 1–19.

73. Muller HJ. 1928. The measurement of gene mutation rate in *Drosophila*, its high variability, and Its dependence upon temperature. Genetics 13: 279–357.

74. Nattestad M, Aboukhalil R, Chin CS, Schatz MC. 2021. Ribbon: intuitive visualization for complex genomic variation. Bioinformatics 37: 413–415.

75. Ness RW, Morgan AD, Colegrave N, Keightley PD. 2012. Estimate of the spontaneous mutation rate in *Chlamydomonas reinhardtii*. Genetics 192: 1447–1454.

76. Ness RW, Morgan AD, Vasanthakrishnan RB, Colegrave N, Keightley PD. 2015. Extensive de novo mutation rate variation between individuals and across the genome of *Chlamydomonas reinhardtii*. Genome Res 25: 1739–1749.

77. O’Donnell S, Fischer G. 2020. MUM&Co: accurate detection of all SV types through whole-genome alignment. Bioinformatics 36: 3242–3243.

78. Ohno S. 1970. Evolution by gene duplication. Allen & Unwin; Springer-Verlag, London,New York,.

79. Parks MM, Lawrence CE, Raphael BJ. 2015. Detecting non-allelic homologous recombination from high-throughput sequencing data. Genome Biol 16: 72.

80. Potter S, Bragg JG, Blom MP, Deakin JE, Kirkpatrick M, Eldridge MD, Moritz C. 2017. Chromosomal speciation in the genomics era: disentangling phylogenetic evolution of rock-wallabies. Front Genet 8: 10.

81. Quinlan AR, Hall IM. 2010. BEDTools: a flexible suite of utilities for comparing genomic features. Bioinformatics 26: 841–842.

82. Rech GE, Radio S, Guirao-Rico S, Aguilera L, Horvath V, Green L, Lindstadt H, Jamilloux V, Quesneville H, Gonzalez J. 2022. Population-scale long-read sequencing uncovers transposable elements associated with gene expression variation and adaptive signatures in *Drosophila*. Nat Commun 13: 1948.

83. Rhoads A, Au KF. 2015. PacBio sequencing and Its applications. Genomics Proteomics Bioinformatics 13: 278–289.

84. Ricci M, Peona V, Guichard E, Taccioli C, Boattini A. 2018. Transposable elements activity is positively related to rate of speciation in mammals. J Mol Evol 86: 303–310.

85. Robinson JT, Thorvaldsdóttir H, Winckler W, Guttman M, Lander ES, Getz G, Mesirov JP. 2011. Integrative Genomics Viewer. Nat Biotechnol 29: 24–26.

86. Ruiz-Ruano FJ, López-León MD, Cabrero J, Camacho JPM. 2016. High-throughput analysis of the satellitome illuminates satellite DNA evolution. Sci Rep 6: 28333.

87. Sedlazeck FJ, Rescheneder P, Smolka M, Fang H, Nattestad M, von Haeseler A, Schatz MC. 2018. Accurate detection of complex structural variations using single-molecule sequencing. Nat Methods 15: 461–468.

88. Sfeir A, Symington LS. 2015. Microhomology-mediated end joining: a back-up survival mechanism or dedicated pathway? Trends Biochem Sci 40: 701–714.

89. Shen W, Le S, Li Y, Hu F. 2016. SeqKit: a cross-platform and ultrafast toolkit for FASTA/Q file manipulation. PLoS One 11: e0163962.

90. Sinha S, Li F, Villarreal D, Shim JH, Yoon S, Myung K, Shim EY, Lee SE. 2017. Microhomology-mediated end joining induces hypermutagenesis at breakpoint junctions. PLoS Genet 13: e1006714.

91. So A, Le Guen T, Lopez BS, Guirouilh-Barbat J. 2017. Genomic rearrangements induced by unscheduled DNA double strand breaks in somatic mammalian cells. Febs J 284: 2324–2344.

92. Song JM, Guan Z, Hu J, Guo C, Yang Z, Wang S, Liu D, Wang B, Lu S, Zhou R et al. 2020. Eight high-quality genomes reveal pan-genome architecture and ecotype differentiation of *Brassica napus*. Nat Plants 6: 34–45.

93. Sui Y, Qi L, Wu JK, Wen XP, Tang XX, Ma ZJ, Wu XC, Zhang K, Kokoska RJ, Zheng DQ et al. 2020. Genome-wide mapping of spontaneous genetic alterations in diploid yeast cells. Proc Natl Acad Sci U S A 117: 28191–28200.

94. Sung W, Ackerman MS, Miller SF, Doak TG, Lynch M. 2012. Drift-barrier hypothesis and mutation-rate evolution. Proc Natl Acad Sci U S A 109: 18488–18492.

95. Tusso S, Suo F, Liang Y, Du LL, Wolf JBW. 2022. Reactivation of transposable elements following hybridization in fission yeast. Genome Res 32: 324–336.

96. Uzunović J, Josephs EB, Stinchcombe JR, Wright SI. 2019. Transposable elements are important contributors to standing variation in gene expression in *Capsella grandiflora*. Mol Biol Evol 36: 1734–1745.

97. van Dijk K, Xu H, Cerutti H. 2006. Epigenetic Silencing of transposons in the green alga Chlamydomonas reinhardtii. In Small RNAs: Analysis and Regulatory Functions, doi:10.1007/978-3-540-28130-6_8 (ed. W Nellen, C Hammann), pp. 159–178. Springer Berlin Heidelberg, Berlin, Heidelberg.

98. van’t Hof AE, Campagne P, Rigden DJ, Yung CJ, Lingley J, Quail MA, Hall N, Darby AC, Saccheri IJ. 2016. The industrial melanism mutation in British peppered moths is a transposable element. Nature 534: 102–105.

99. Weischenfeldt J, Symmons O, Spitz F, Korbel JO. 2013. Phenotypic impact of genomic structural variation: insights from and for human disease. Nat Rev Genet 14: 125–138.

100. Weissensteiner MH, Bunikis I, Catalan A, Francoijs KJ, Knief U, Heim W, Peona V, Pophaly SD, Sedlazeck FJ, Suh A et al. 2020. Discovery and population genomics of structural variation in a songbird genus. Nat Commun 11: 3403.

101. Weissman JL, Fagan WF, Johnson PLF. 2019. Linking high GC content to the repair of double strand breaks in prokaryotic genomes. PLoS Genet 15: e1008493.

102. Yoder AD, Tiley GP. 2021. The challenge and promise of estimating the de novo mutation rate from whole-genome comparisons among closely related individuals. Mol Ecol 30: 6087–6100.

103. Yun T, Li H, Chang PC, Lin MF, Carroll A, McLean CY. 2020. Accurate, scalable cohort variant calls using DeepVariant and GLnexus. Bioinformatics doi:10.1093/bioinformatics/btaa1081.

104. Zhang C, Wu-Scharf D, Jeong BR, Cerutti H. 2002. A WD40-repeat containing protein, similar to a fungal co-repressor, is required for transcriptional gene silencing in *Chlamydomonas*. Plant J 31: 25–36.

105. Zhang J, Yu C, Pulletikurti V, Lamb J, Danilova T, Weber DF, Birchler J, Peterson T. 2009. Alternative *Ac*/*Ds* transposition induces major chromosomal rearrangements in maize. Genes Dev 23: 755–765.

106. Zhang L, Chaturvedi S, Nice CC, Lucas LK, Gompert Z. 2022. Population genomic evidence of selection on structural variants in a natural hybrid zone. Mol Ecol doi:10.1111/mec.16469.

107. Zhang X, Zhao M, McCarty DR, Lisch D. 2020. Transposable elements employ distinct integration strategies with respect to transcriptional landscapes in eukaryotic genomes. Nucleic Acids Res 48: 6685–6698.

108. Zhao H, Zhang W, Chen L, Wang L, Marand AP, Wu Y, Jiang J. 2018. Proliferation of regulatory DNA elements derived from transposable elements in the maize genome. Plant Physiol 176: 2789–2803.

