## Supplementary Material for "Rates and spectra of *de novo* structural mutation in *Chlamydomonas reinhardtii*"


**Appendix S1.** Ancestor genome assembly and genome-wide callability

- **Figure S1.** Assembly metrics
- **Figure S2.** Length of the callable genome

**Appendix S2.** Rejected structural mutation calls

- **Figure S3.** Number of variants obtained with different variant calling methods
- **Figure S4.** Distance of variants to major repeats

**Appendix S3.** Genomic distribution of structural mutations

- **Figure S5.** SM genomic distribution

**Appendix S4.** CC-2931 translocations and inversions

- **Figure S6.** Translocations and inversions in CC-2931 MA lines

**Appendix S5.** *Dualen-4b_cRei* target site duplications

- **Figure S7.** *Dualen-4b_cRei* target site duplications (TSD) in CC-1952

**Appendix S6.** Supporting material for homology-mediated structural mutations

- **Figure S8.** Sequences putatively involved in macrohomology and NAHR
- **Figure S9.** A putative MMEJ-mediated deletion
- **Figure S10.** Sequences putatively involved in microhomology and MMEJ

**Appendix S7.** Examples of structural mutation visualisation

- **Figure S11-S25.** SM snapshot examples

**References**

**Appendix S1.** Ancestor genome assembly and genome-wide callability

#### *De novo assembly*

Chromosome-level ancestor assemblies were produced using a hybrid approach. The CC-2931 ancestor CLR reads were assembled independently with wtdbg2 (Ruan and Li, 2020) with the parameters "-g 111m -x sq" and Flye v2.8.2 (Kolmogorov et al. 2019) with the parameter "-g 111.1m". The Flye assembly was further processed using purge_dups v1.2.5 (Guan et al. 2020). The resulting contig-level assemblies were error corrected using PacBio reads and two iterations of the Arrow module of GCpp (https://github.com/PacificBiosciences/gcpp). Further polishing was performed using Illumina reads and one iteration of Pilon v2.14 ("--fix bases) (Walker et al. 2014), with post-processing to restrict polishing to the correction of single nucleotide changes and INDELs of 5 bp or less (see Craig et al. 2021). Illumina data from all 14 CC-2931 MA lines (Ness et al. 2015) were used after subsampling each MA line readset to 10% coverage to ensure that no short mutations, which are expected to be unique to MA lines, were incorporated by polishing. Both polished assemblies were then mapped to the near-complete chromosome-level assembly of *C. reinhardtii* CC-1690 (O'Donnell et al. 2020) using minimap2 v2.17 (Li 2018). Scaffolding was manually performed to produce chromosomes based on inspection of the resulting pairwise alignment format (PAF) files. The Flye assembly was given precedence, and where possible the wtdbg2 assembly was used to fill gaps between Flye contigs, extend Flye contigs at chromosome termini, or to fix misassemblies. All steps that involved merging Flye and wtdbg2 contigs were manually checked against the raw reads using IGV 2.4.2 (Robinson et al. 2011). Blocks of 10,000 Ns were added to gaps that could not be filled and to terminal contigs that did not end in telomeric repeats. Finally, a circular plastome assembly was produced using Circlator (Hunt et al. 2015). Putative plastome reads were extracted by mapping against the existing plastome assembly of strain CC-503 (Gallaher et al. 2018) and a preliminary assembly was produced using Canu v2.1.1 ("genomeSize=0.2m") (Koren et al. 2017). The single resulting contig was then passed to Circlator together with the error-corrected reads output by Canu for circularisation. The CC-2931 mitogenome was previously assembled by Smith & Lee (2008) and was directly appended.

The CC-1952 ancestor was assembled in a similar manner as described above, except that HiFi reads were assembled using Flye and hifiasm v0.15.3 ("-l0 -f0") (Cheng et al. 2021). The hifiasm contigs were given precedence during scaffolding given the discovery of high error rates in HiFi-based Flye assemblies at interspersed repeats (see below), with Flye contigs only used for gap filling and contig extension. No error correction was performed. Plastome assembly was performed as above, except the preliminary assembly was produced using HiCanu v2.1.1 (Nurk et al. 2020). The mitogenome was appended from Smith & Lee (2008).

#### *Assembly quality and callability*

Assembly contiguity was high for both ancestor and mutation accumulation (MA) line assemblies, all exhibiting contig-level N50 values >1 Mb (where N50 is the shortest contig length of the minimum set of contigs representing 50% of the total assembly length). Contiguity was higher for the samples sequenced using PacBio CLR, compared to CCS

(HiFi) (Fig. S1A). Genome completeness measured via benchmarking universal single-copy orthologs (BUSCOs; Manni et al. 2021) yielded ≥99% completeness (Fig. S1B).

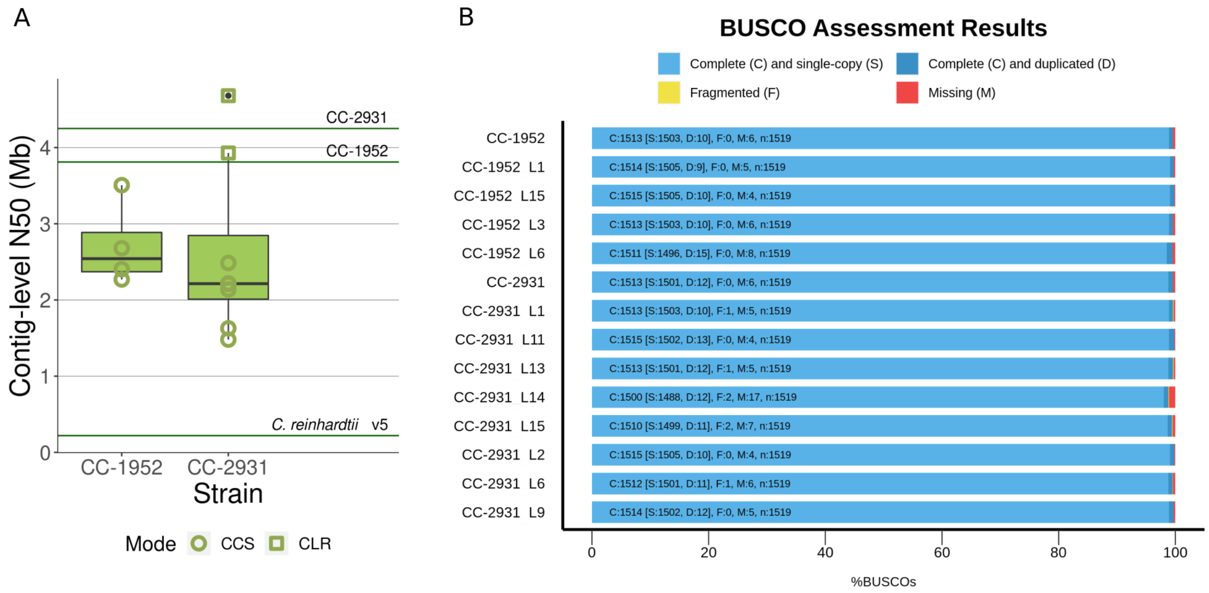


Fig. S1. Assembly metrics. A) Genome contiguity. Contig-level N50 values (in Mb) are shown for CC-1952 and CC-2931 MA line assemblies. The shape of the points identifies the sequencing mode used, PacBio CCS (HiFi, in circles) or CLR (in squares). The horizontal lines show the contiguity for the ancestor assemblies of each strain and the *C. reinhardtii* v5 reference assembly (Blaby et al. 2014). The CC-2931 ancestor assembly was based on PacBio CLR and the CC-1952 ancestor assembly was based on PacBio CCS. B) Genome completeness. Percentage of complete (single copy in light blue, duplicated in dark blue), fragmented (yellow) and missing (red) BUSCOs (chlorophyta_odb10 dataset) in the CC-1952 and CC-2931 ancestors and their MA lines.

We defined nearly 98% of the genome as callable, i.e. of sufficient quality to call structural mutations (SMs). Our definition of the callable genome was based on the ability of genomic regions to align 1) in ancestor self-to-self alignments, and 2) in alignments of all MA lines against the ancestor (allowing for no more than one unmapped MA line assembly, see Methods). Cumulatively, only 1.8 Mb (CC-1952) and 2.3 Mb (CC-2931) of the ancestor genomes could not be assessed for mutations using our long-read datasets. By chromosome, the length of callable sequence was strongly correlated between long-read (PacBio CLR or CCS) and short-read (Illumina) datasets (Pearson's product-moment correlation; $r \geq 0.98$, $P \leq 1.2 \times 10^{-12}$), although callability was on average 27% higher with PacBio. The difference in per-chromosome callability for CC-1952 and CC-2931 between PacBio and Illumina is represented in Fig. S2.

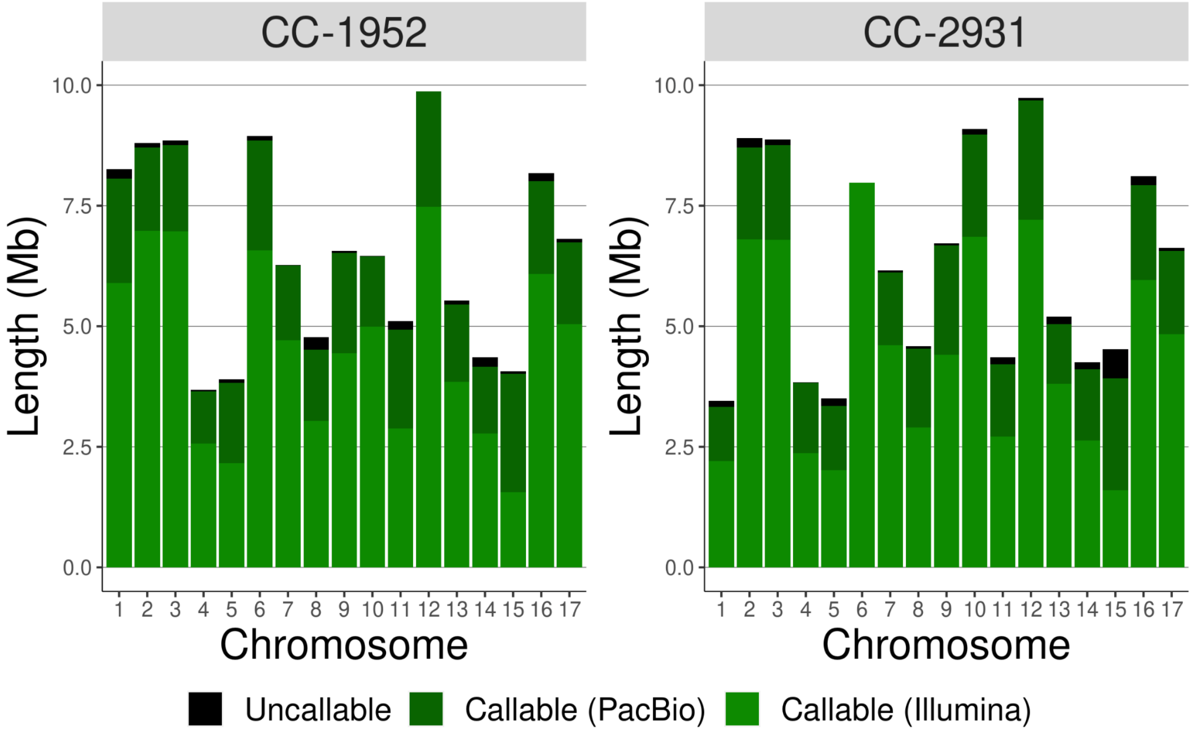


Fig. S2. Genome-wide callability. Bar heights represent total chromosome length for each chromosome (Mb). Colours represent the total sequence that was callable using Illumina data (light green), the increase in callable sequence achieved with PacBio (dark green) and the remaining uncallable sequence (black).

Note that the chromosome lengths differ between the two strains due to chromosomal rearrangements that occurred in the CC-2931 ancestor affecting chromosomes 1, 6 and 10 (see main text). Also note that Illumina-based callability was obtained by aligning reads to the v5 reference genome in our previous study (Ness et al. 2015). Since the v5 reference is based on a strain (CC-503) that is divergent from the MA ancestors, we can not solely attribute the increase in callability achieved with PacBio to differences between sequencing platforms.

### Appendix S2. Rejected structural mutation calls

To obtain an estimate of the performance of structural variant callers, we classified calls in four categories. Variants accepted as *de novo* SMs were classified as confirmed variants (CV). Variants rejected were classified as 'uncallable' if they did not fall within the callable genome (see above). Other rejected variants at callable sites were either classified as unsupported variants (UV) or assembly errors (AE). Both UV and AE calls were not supported by read mapping, but in contrast to UVs, AEs were present in the relevant MA line assembly, so that the calling failure could be attributed to an assembly error rather than to the variant caller. We followed a lenient criterion, including only as UVs the calls that could not be associated with any variant observed in the assembly alignment (even if the variant differed in the type of SM called). Only calls at unique coordinates are considered here.

The total number of variants called was higher for the methods based on assembly alignments than for the read-based Sniffles (Sedlazeck et al. 2018) (Fig. S3). For the two assembly-based methods, vg (Garrison et al. 2018) more than doubled the total number of calls compared to MUM&Co (O'Donell and Fischer 2020). vg was also the method leading to the highest number of CVs (see Fig. 1 in main text and Fig. S3). More variants were rejected for the assembly-based methods than for Sniffles, partly due to the presence of assembly errors. Among the rejected variants whose type could be directly determined by the caller, the majority were insertion (41.9%) and deletion (30.6%) calls, followed by translocations (16.0%).

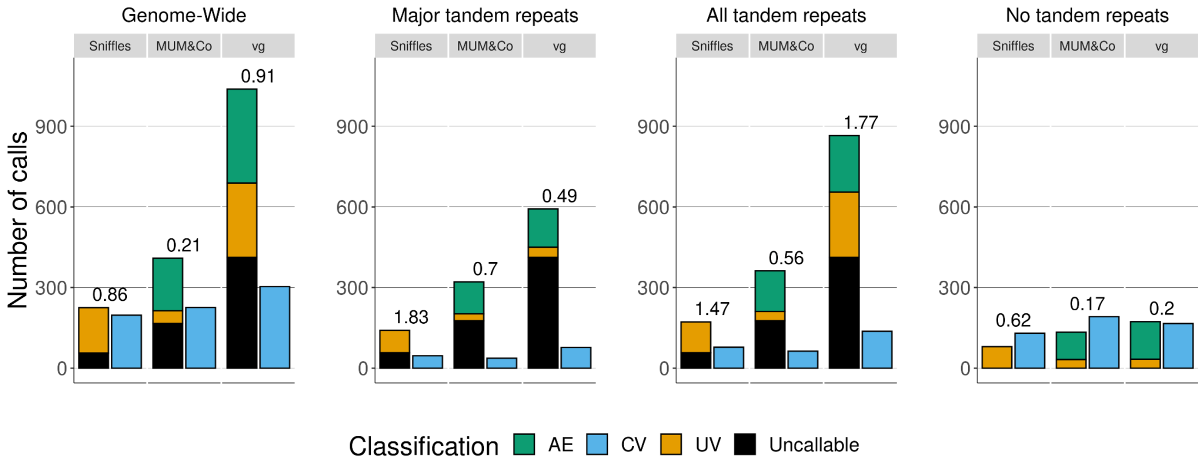


Fig. S3. Number of variants obtained with different variant calling methods at unique genomic coordinates across the eight CC-2931 MA lines. Bars are stacked for accepted (CV: confirmed variant) and rejected calls (UV: unsupported variant, AE: assembly error, and uncallable variant). Values above the bars indicate the ratio of UV over CV. The left-most panel shows the total number of variant calls obtained genome-wide. The second and third panels show calls restricted to those with at least one breakpoint overlapping certain repetitive sequences, either major tandem repeats (centromeres, subtelomeres, ribosomal DNA, satellite DNA >1 kb, and uncallable sites) or all tandem repeats (grouping major tandem repeats with shorter satellites and microsatellites), respectively. The right-most panel shows variant calls outside tandem repeat annotation.

As mentioned in the main text, all three of the approaches returned more rejected calls than confirmed SMs, and rejected call rates were substantially affected by genomic repetitiveness. Using the ratio of genome-wide rejected calls to confirmed SMs in CC-2931 as a benchmark, Sniffles performed best (1.14), followed by MUM&Co (1.81) and vg (3.42). Calls made in uncallable regions (see above) were automatically rejected, since these regions generally consist of major tandem repeats that exceed the length of the PacBio reads, making both genome assembly and variant calling unreliable. MUM&Co and vg returned more variants in uncallable regions than Sniffles (~30% of all MUM&Co and vg variants vs ~15% of Sniffles variants), partly because contig ends in these highly repetitive regions were often called as variants. In callable regions, the proportion of variants rejected was similar across methods (ranging from 46.2% to 67.4%). Tandem repeats of all lengths were a major source UVs, and to a lesser extent AEs (Fig. S3). This was especially true for vg, with ~90% of UVs found in tandem repeats. Notably, this error rate was not restricted to the most complex tandem repeats (e.g. centromeres and major satellites), but was in fact most pronounced in microsatellites. This was generally due to multiple variants being called within the same repeat that cancelled each other out (e.g. an insertion and deletion of similar lengths within a single microsatellite). When excluding all tandem repeats, the ratio of UVs to confirmed SMs improved considerably for MUM&CO (0.21 to 0.17), Sniffles (0.86 to 0.62) and vg (0.91 to 0.20).

For the assembly-based methods, the majority of variants rejected at callable sites were AEs (41.8% MUM&CO, 37.7% vg). This result highlights the importance of the assembly process. Here we chose Flye (Kolmogorov et al. 2019) as an assembler since it produced the best haploid approximation of the *C. reinhardtii* genome (see Methods). However, we later noticed that Flye introduced many errors at interspersed repeats. Flye uses secondary alignment to error-correct repetitive regions, which apparently resulted in variants present in other copies of a given interspersed repeat being sporadically incorporated (e.g. at a TE sequence on chromosome 1, variants found in another copy of that TE elsewhere in the genome could be assembled). These assembly errors predominantly introduced short variants, although several AEs were also associated with this issue. Although this did not affect our results following our curation of all mutations, we caution that Flye is likely to increase the rate of assembly errors at interspersed repeats. Finally, other types of AEs were observed that were not necessarily specific to Flye. We frequently observed cases where the incorrect number of monomers were assembled in satellite DNA (which were subsequently called as expansions or contractions but were unsupported by read mapping). Furthermore, as mentioned above, both MUM&CO and vg regularly called the end of contigs as variants e.g. as deletions relative to the ancestor assembly. These AEs could easily be removed by removing variant calls associated with contig ends, which approximately halved the number of AEs remaining after removing tandem repeats for both MUM&Co and vg.

Finally, to further explore the relationship between genome repetitiveness and rejected variants, we estimated the distance from different types of calls to major tandem repeats, including centromeres, subtelomeres and satellite DNA >1 kb (Fig. S4). The density of CVs was significantly higher in regions further from these major repeats than for rejected calls, and the opposite trend was observed for rejected variants (Wilcoxon rank sum test, W test, $P < 2.2 \times 10^{-16}$). Still, a peak of CVs was observed in these repetitive annotations. This is because many variants could be called and confirmed in these highly mutable regions (e.g. contractions and expansions) despite their repetitiveness. The peak observed for uncallable variants at a distance longer than 1 kb from a major repeat was mainly due to regions close to gaps in the ancestor assemblies that we failed to identify as tandemly repeated (see Methods), but were nonetheless obviously highly complex and repetitive when viewed as dotplots. A small proportion of uncallable regions were also associated with TE-rich regions, which, for example, are particularly prominent on chromosome 15 (see Fig. S2).

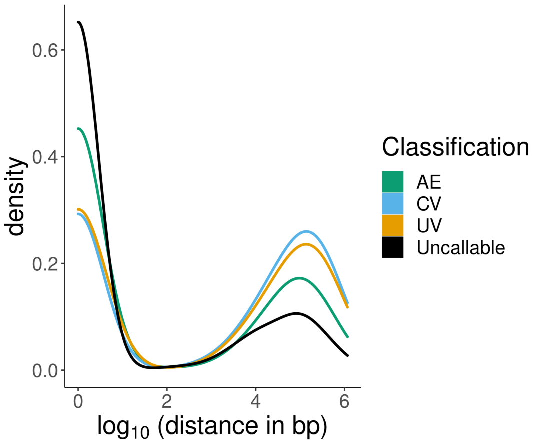


Fig. S4. Distance of variants to major repeats (in base pairs; $log_{10}$ scale). The plot shows the density of different classifications of calls (in colours; CV: confirmed variant, UV: unsupported variant, AE: assembly error, and uncallable variant) with their distance to major repeats (satellite DNA >1 kb, ribosomal DNA, centromeres and subtelomeric sequences). Data comes from 8 MA lines with CC-2931 genotype.

**Appendix S3.** Distribution of structural mutations

To explore the distribution of SMs across the genome, and in particular their distribution relative to functional sequences, we generated ~20 Gb of RNA-seq data for the CC-2931 ancestor and annotated coding sequences, introns and untranslated regions (UTRs). We also divided intergenic regions into "proximal" (within 500 bp of a gene start or end) and "distal" sequences (>500 bp from a gene), the latter category largely capturing large and highly repetitive intergenic regions in the compact *C. reinhardtii* genome (including the centromeres and subtelomeres).

We then compared the observed distribution of SM coordinates to the expectation based on random sampling of the callable genome. Such an analysis required a precise definition of SM coordinates. While mobile insertions could be assigned to a single insertion site (see main text), we defined the coordinates of SMs that affected a length of sequence (i.e. duplications, deletions and inversions) in two ways. First, we considered the entire length span of each SM. Second, we considered both the start and end coordinates of such SMs. While the former approach takes into account the wider possible effects of mutation (e.g. the duplication of entire genes), the latter explores whether the exact breakpoints of SMs were enriched at any particular genomics features. We did not analyse contractions, expansions and mobile excisions.

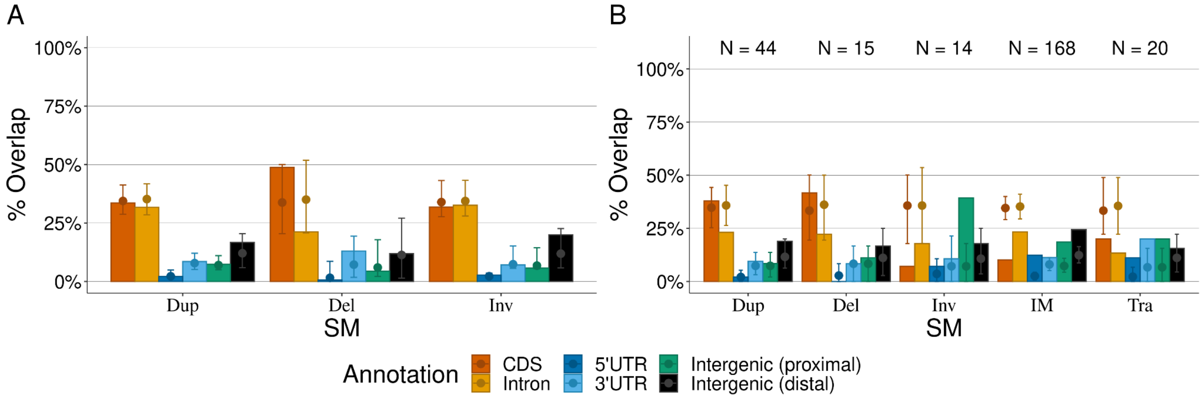


Fig. S5. SM genomic distribution. A) Overlap of SMs with genome annotations. The proportion of overlapping sequence is shown in bars for different functional annotations (in colours) and types of SM. The expectation for SMs of the same length randomly distributed along the callable genome is shown with a point (median overlap) and error bars (95 % confidence interval). B) Distribution of SM flanking coordinates (start and end) relative to genomic annotations. The proportion of sequence overlap of different SM types is shown for each functional annotation class (in colours). The random expectation is shown by a point (median proportion of sequence overlap) and error bars (95 % confidence interval). Expected values are based on 1,000 samples. SM types included in A) and B) are: Duplications (Dup), deletions (Del), inversions (Inv), insertion of mobile elements (IM) and translocations (Tra). The total number of these SMs is shown with numbers on the top of B).

The span of duplications, deletions and inversions intersected with genomic annotations as expected under random distribution (Fig. S5A). This included their intersection with coding sequence, suggesting that many of these SMs could have large fitness effects. Furthermore, the breakpoints of duplications and deletions were also distributed randomly (Fig. 5SB). The breakpoints of inversions and translocations did exhibit a bias towards UTRs and gene-proximal intergenic sequence (Fig. S5B), a result that we attributed to their association with TEs (see main text). These results support a lack of selection in our MA experiment, as expected due to the regular population bottlenecks.

**Appendix S4.** CC-2931 translocations and inversions

In most CC-2931 MA lines, we observed reciprocal and non-reciprocal translocations (in 5 MA lines out of 8). The most remarkable example of genome reorganisation is provided by CC-2931 MA line L13, where we observed a total of 8 translocations, 4 of which were non-reciprocal (Fig. S6A). Reciprocal translocations included those between chromosomes 6 and 8, 7 and 11, 10 and 12, and between 9 and 16 (see Fig. 4C). These translocation events included events that were likely mediated by *Un3*/*Un12* TE insertions (e.g. chromosomes 7 and 11), but also *CryptonF-1_cRei* insertions (chromosomes 6 and 8). The remaining events were non-reciprocal and involved chromosomes 3, 7, 8 and 10. These events were mediated by *Un3* and *Un12* insertions, which presumably inserted simultaneously on the involved chromosomes.

We observed four reciprocal translocations in line 15 (Fig. S6B). However, only one of these events, between chromosomes 1 and 11, was TE-mediated (by *CryptonF-1_cRei*). Finally, although there were no translocations in line 14, we observed four inversions (Fig. S6C, see main text Fig. 4C for the fourth inversion on chromosome 16). Three of these four inversions were apparently mediated by *CryptonF-1_cRei*, involving an insertion or excision at one breakpoint and a 'CAYCG' target site at the other.

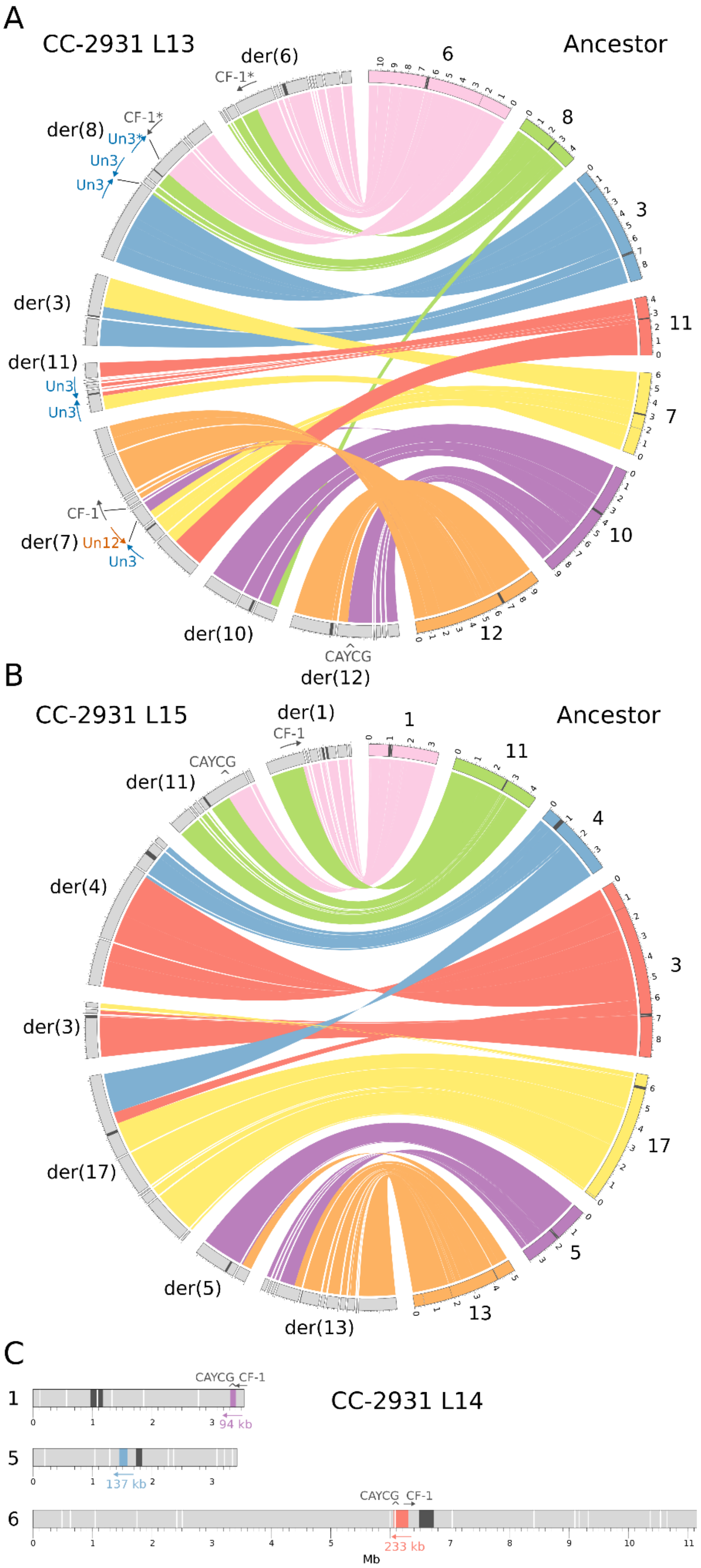
A
CC-2931 L13
Ancestor
der(6)
CF-1*
der(8)
Un3*
Un3
der(3)
der(11)
der(7)
Un12
CF-1
der(10)
der(12)
CAYCG
6
8
3
11
7
10
12
B
CC-2931 L15
Ancestor
der(1)
CF-1
CAYCG
der(11)
der(4)
der(3)
der(17)
der(5)
der(13)
1
11
4
3
17
5
13
C
CC-2931 L14
CAYCG
CF-1
94 kb
137 kb
233 kb
Mb

Fig. S6. Translocations and inversions in CC-2931 MA lines. Translocations in CC-2931 MA lines 13 (A) and 15 (B). The ancestor chromosomes are represented on the right half of each Circos plot, while the MA line chromosomes (as contigs) are on the left. Chromosome numbers are given for the ancestor, while derived chromosomes (denoted with "der") are provided for the MA lines based on centromere annotation (dark grey regions). Scale is in megabases. CF-1 indicates *CryptonF-1_cRei* insertions. The direction of the arrows indicates the 5' to 3' orientation of the TE sequence. An asterisk next to a TE name indicates that the insertion is incomplete (observed at one pair of repair sites, where a *CryptonF-1_cRei* insertion was fragmented and fused with an incomplete *Un3* element). C) Inversions in CC-2931 MA line 14. The three coloured regions depict inverted regions, while dark grey regions are centromeres and white regions are assembly gaps.

### **Appendix S5.** *Dualen-4b_cRei* target site duplications

Target site duplications associated with LINEs are caused by resolution of the DNA nicks introduced during insertion, with the distance between cleavage sites corresponding to TSD length. *Dualen* are unique among LINEs in that they encode both restriction-like endonuclease and apurinic/apyrimidinic endonuclease-like endonuclease (Kojima and Fujiwara 2005). It is possible that the dual activity of endonucleases is responsible for the giant target site duplications observed here. Interestingly, we observed some large duplications flanking other families of *Dualen* elements in the ancestor assemblies (i.e. TEs that were not active in the MA experiment), suggesting that this phenomenon is not specific to *Dualen-4b_cRei* and may generally be associated with *C. reinhardtii Dualen* elements.

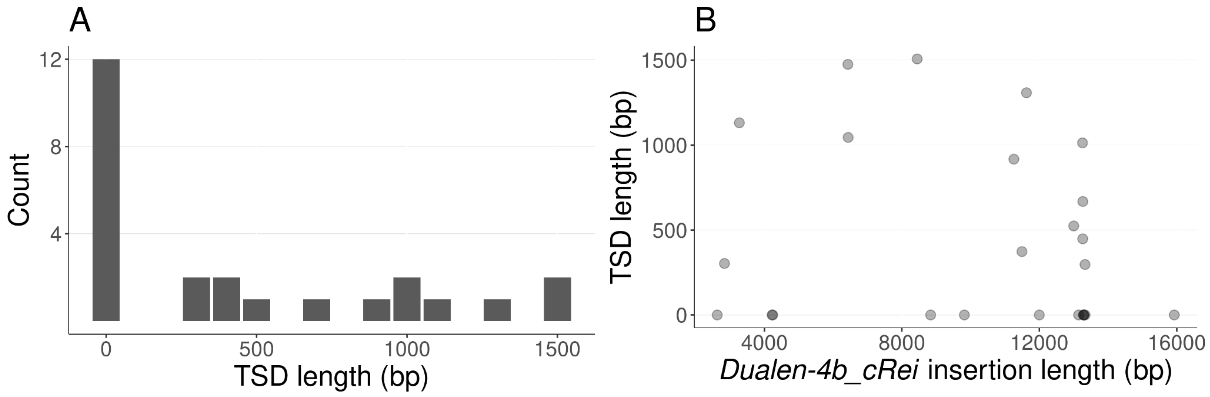


Fig. S7. *Dualen-4b_cRei* target site duplications (TSD) in CC-1952. A) Histogram of TSD lengths (in base pairs). For simplicity, *Dualen-4b_cRei* insertions without associated duplications are assumed to have TSDs of length zero . B) Distribution of *Dualen-4b_cRei* insertions and their corresponding TSD lengths. Darker points indicate a higher count of inserted elements.

Out of the 25 *Dualen-4b_cRei* insertions detected in CC-1952, 13 (52%) were associated with target site duplications long enough to be classified as SMs (median length = 915, 95% CI = [298.8, 1497.4], see Fig. S7A). The complete *Dualen-4b_cRei* element is nearly 13 kb in length. 40% of *Dualen-4b_cRei* insertions were complete, with the remainder being 5' truncated. There was no apparent relation between the length of the insertion and the length of the target site duplication (Pearson's product-moment correlation; $t_{23} = -0.81$, $r = -0.17$, $P = 0.43$, Fig. S7B).

**Appendix S6.** Supporting material for homology-mediated structural mutations

As mentioned in the main text, we used methods similar to Belyeu et al. (2021) to investigate the association of SM breakpoints with patterns of homology that could potentially associate SMs with different double-strand break (DSB) repair pathways. Figure S8 shows the alignment of the flanking sequence of three duplications, supporting the involvement of non-allelic homologous recombination (NAHR) in these events. Figure S9 shows alignment of 20 bp sequences putatively involved in MMEJ repair, and explaining a total of 5 deletions. Figure S10 shows the remarkable case of a deletion attributed to microhomology-mediated end-joining (MMEJ) that was flanked by a substantial number of single nucleotide mutations and INDELs, potentially due to the high mutagenicity of MMEJ (Sinha et al. 2017).

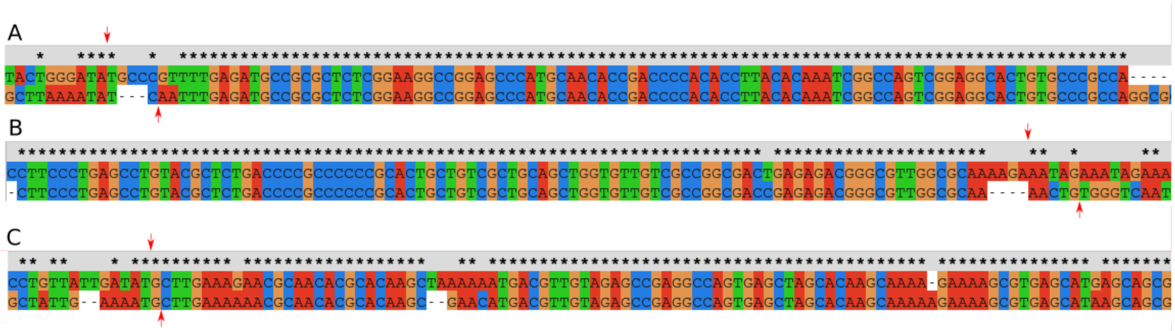


Fig. S8. Sequences putatively involved in macrohomology and NAHR. Visualisation and alignment made with ClustalX (Larkin et al. 2007), using sequences extending 100 bp upstream and downstream from the start (top) and end (bottom) coordinates of the duplication. Breakpoints are indicated with red arrows. Duplications shown are: In CC-1952, chromosome_11:3515460-3516199 (L6); in CC-2931, B) chromosome_06:6195074-6198035 (L13), and C) chromosome_15:2924211-2939343 (L2).

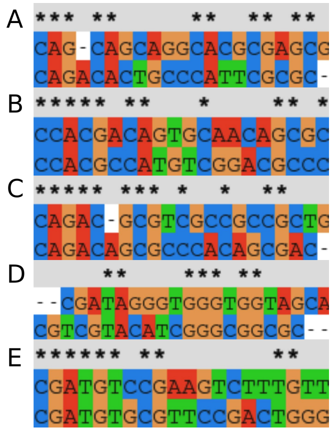


Fig. S9. Sequences putatively involved in microhomology and MMEJ. Visualisation and alignment made with ClustalX (Larkin et al. 2007), from 20 bp sequences downstream from start and end coordinates (5'→ 3' sense) of the following deletions: In CC-1952, chromosome_01:7563718-7563872 (L6); in CC-2931, B) chromosome_02:7522319-7522396 (L1), C) chromosome_03:227504-227725 (L2), D), chromosome_10:7918878-7923936 (L15), and E) chromosome_12:2502394-2502708 (L11).

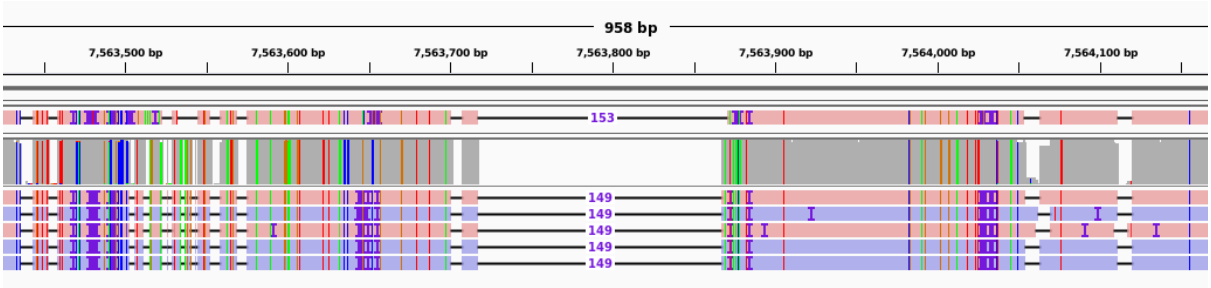


Fig. S10. A putative MMEJ-mediated deletion. IGV visualisation of deletion in CC-1952 MA line 6 at chromosome_01:7563718-7563872, surrounded by multiple SNMs and INDELs.

**Appendix S7.** Examples of structural mutation visualisation

All accepted SMs were manually inspected using the Integrative Genomics Viewer (IGV, Robinson et al. 2011). IGV snapshots show both assembly-to-reference and read-to-reference alignments, with assembly alignments on the top track and read alignments following below (e.g. see assembly with deletion in the top track of Fig. S11, and reads supporting the deletion aligned below). For clarity, most alignments don't show soft-clipped bases, although viewing alignments with soft-clipping turned on was often highly informative. Some complex SMs, particularly rearrangements, were better visualised with Ribbon (Nattestad et al. 2021), which provides a view of the full extent of read mapping across the genome (e.g. translocations, see below), including a single-read view panel. Only read alignment files were used with Ribbon. Both IGV and Ribbon use colours to differentiate forward and reverse mapping reads. Rearrangements much larger than the reads were also visualised from pairwise alignment format (PAF) files generated by minimap2 (Li 2018). Entries in the PAF file were filtered to retain regions with mapping quality of 60 (i.e. uniquely mapping regions). For simplicity, only reads from the mutated MA line are shown.

*Deletions*

Deletions of all lengths were usually clearly visualised in IGV, both in the assembly and read tracks (see Fig. S11).

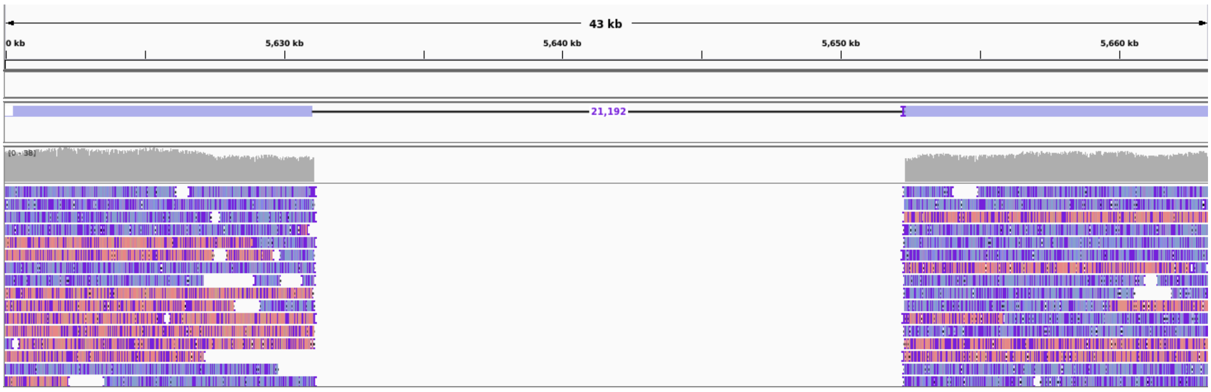


Fig. S11. SM snapshot example: deletion (chromosome_16:5631094-5652288; CC-2931 MA line 2). IGV visualisation.

Considered as deletions, we observed two examples of chromosome arm truncations followed by telomere healing at an internal site (see main text). Fig. S12 shows one of these events in CC-1952 MA line 15, where the telomeric repeat ('TTTTAGGG' in *C. reinhardtii*) can be observed as soft-clipped bases starting ~94 kb from the ancestral telomere on the right arm of chromosome 11.

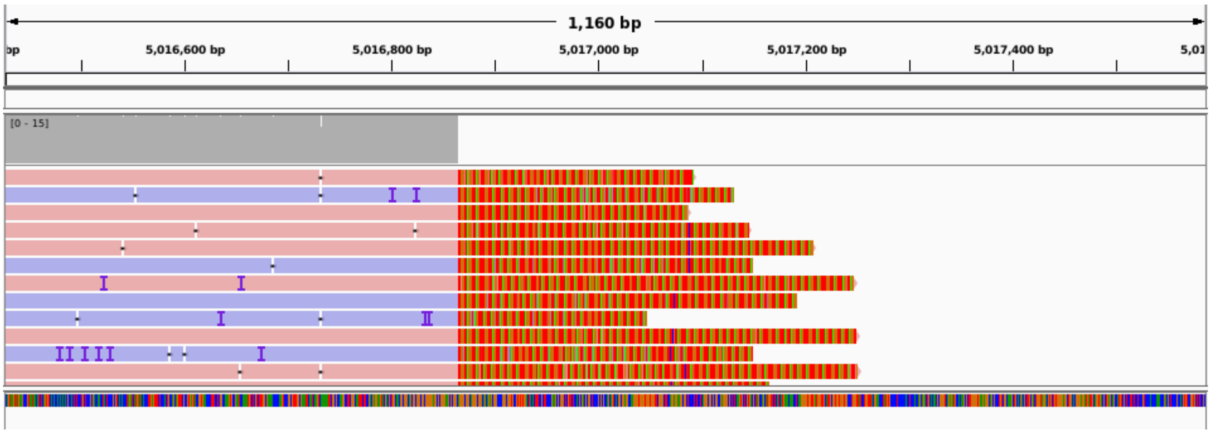


Fig. S12. SM snapshot example: deletion (chromosome_11:5016865-5111318; CC-1952 MA line 15). IGV visualisation (showing soft-clipped bases).

*Contractions and Expansions*

Deletions and insertions of tandemly repeated monomers were annotated as contractions and expansions, respectively. The deleted or inserted segments were often not consecutively aligned among reads, but could clearly be found scattered across the entire repeat annotation (Figs. S13 and S14). In the case of some large unannotated satellites (i.e. those with lengths that exceed the detection limit of Tandem Repeats Finder), tandemly repeated monomers were determined using blastn (Camacho et al. 2009) or dot-plot visualisation with MAFFT (Katoh et al. 2019).

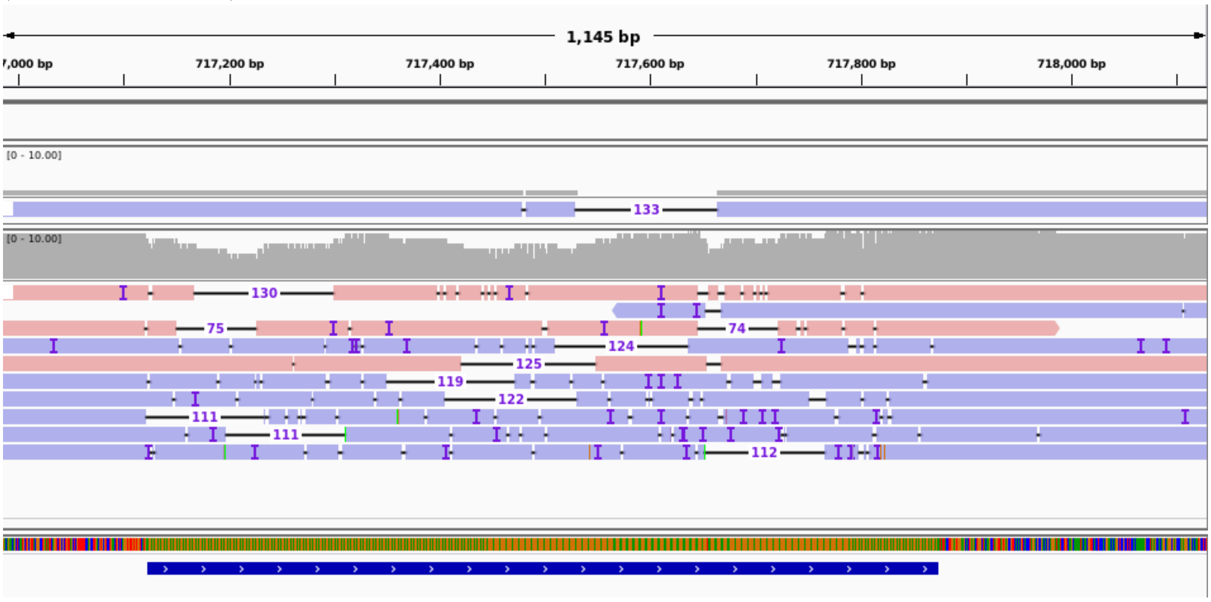


Fig. S13. SM snapshot example: contraction (chromosome_01:717531-717664; CC-2931 MA line 6). The bottom blue line indicates a microsatellite (AGGGAG) annotation. IGV visualisation.

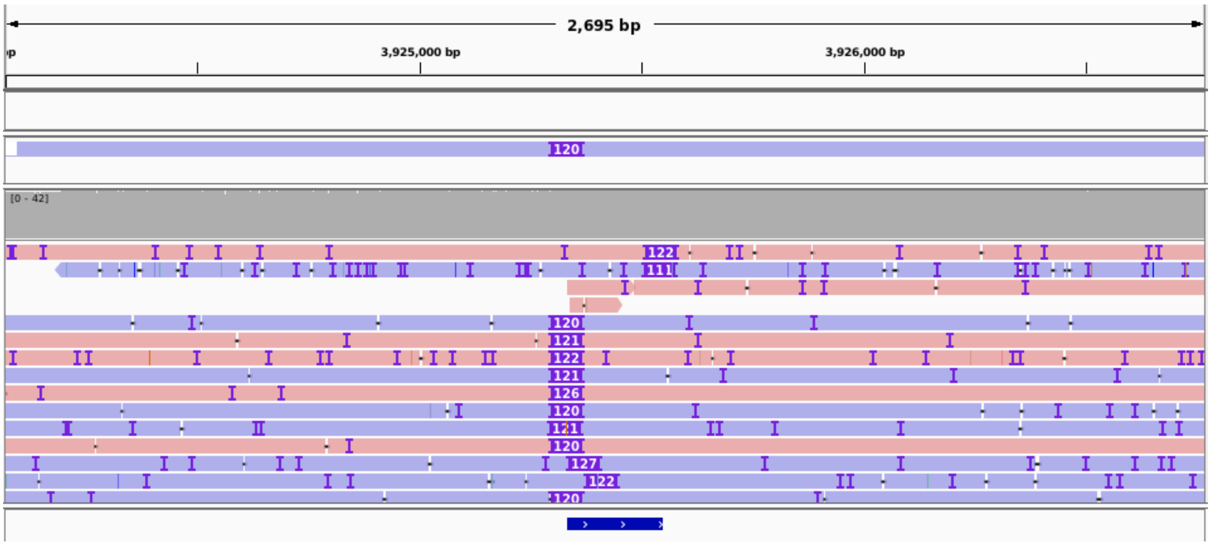


Fig. S14. SM snapshot example: expansion (chromosome_16:3925336-3925337; CC-2931 MA line 1). The bottom blue line indicates a microsatellite (TGTGCG) annotation. IGV visualisation.

*Duplications*

When duplications were shorter than the reads (~25 kb), they were clearly visualised in IGV, both in assembly and read alignments (Fig. S15). However, large duplications were usually collapsed (i.e. absent) from the genome assemblies and could only be called from the read alignments. These were also visualised with Ribbon (Fig. S16).

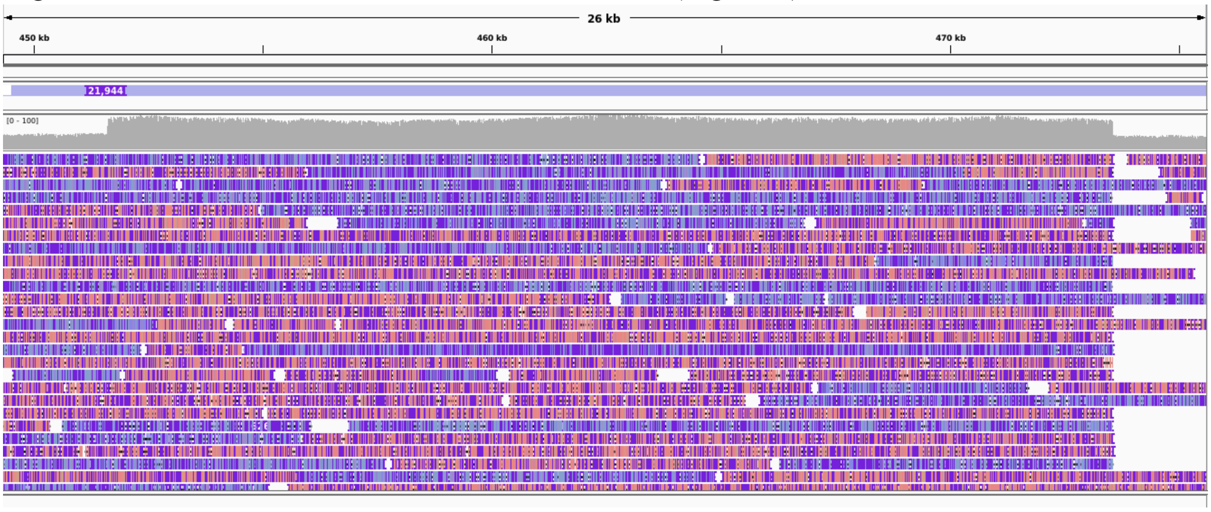


Fig. S15. SM snapshot example: duplication (chromosome_12:451612-473558; CC-2931 MA line 9). IGV visualisation.

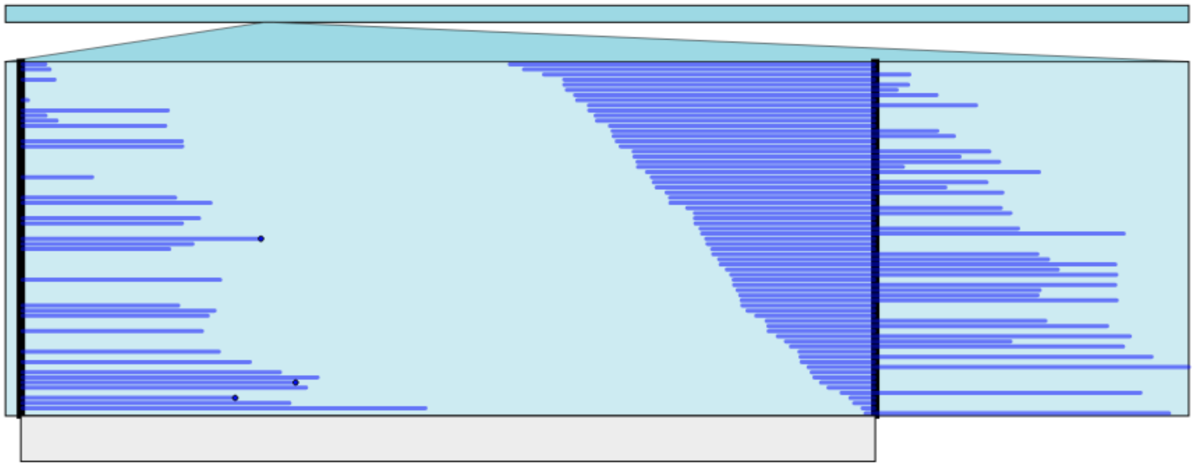

Fig. S16. SM snapshot example: duplication (chromosome_06:2384024-2444969; CC-2931 MA line 14). Ribbon visualisation. Each line is a read. Notice that ~50% of reads map across the right breakpoint (thick black line) while the remaining half are split and map back to the left breakpoint, a clear characteristic of a duplication.

Duplications and deletions could also occur at genomic regions containing some degree of repetitiveness. In principle, duplications in tandemly repeated regions were annotated as expansions (and deletions as contractions, see Methods). However, if the duplication was larger than the span of the tandemly repeated sequence (i.e. if it included newly duplicated sequence), we retained its annotation as duplication. The same was true for deletions that only partly overlapped tandemly repeated sequences. One example is shown in Fig. S17, where a duplication in CC-2931 MA line 13 exceeds the tandemly repeated sequence in the ancestor. In fact, one possible mechanism explaining duplication here might be non-allelic homologous recombination (see Appendix S6).

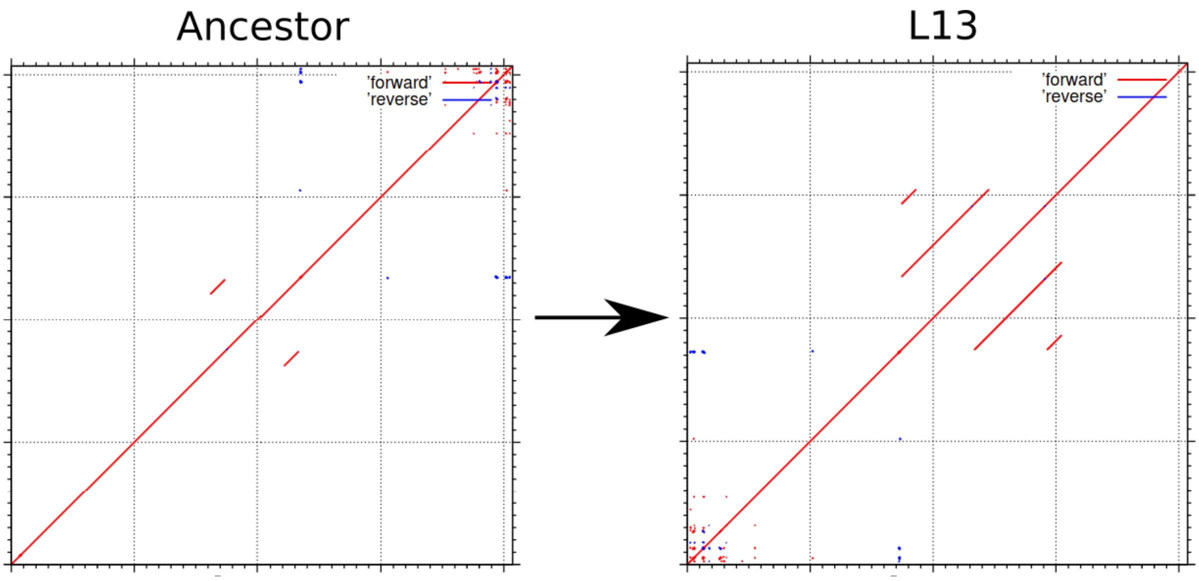


Fig. S17. SM snapshot example: duplication (chromosome_06:6195074-6198035; CC-2931 MA line 13). MAFFT visualisation.

*Inversions*

Similar to duplications, inverted regions and their breakpoints were usually clearly visualised in IGV and Ribbon (Figs. S18, S19). Some extremely large inversions were directly detectable from minimap2 PAF files (see ~80 kb example from a PAF file in Fig. S20).

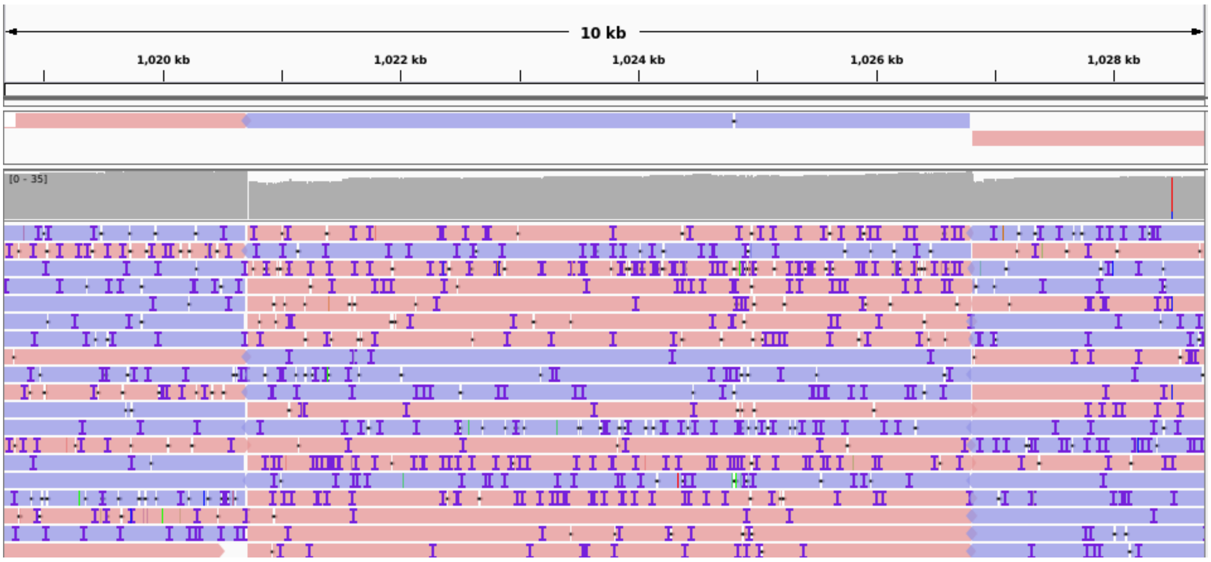


Fig. S18. SM snapshot example: inversion (chromosome_06:1020714-1026818; CC-2931 MA line 15). IGV visualisation.

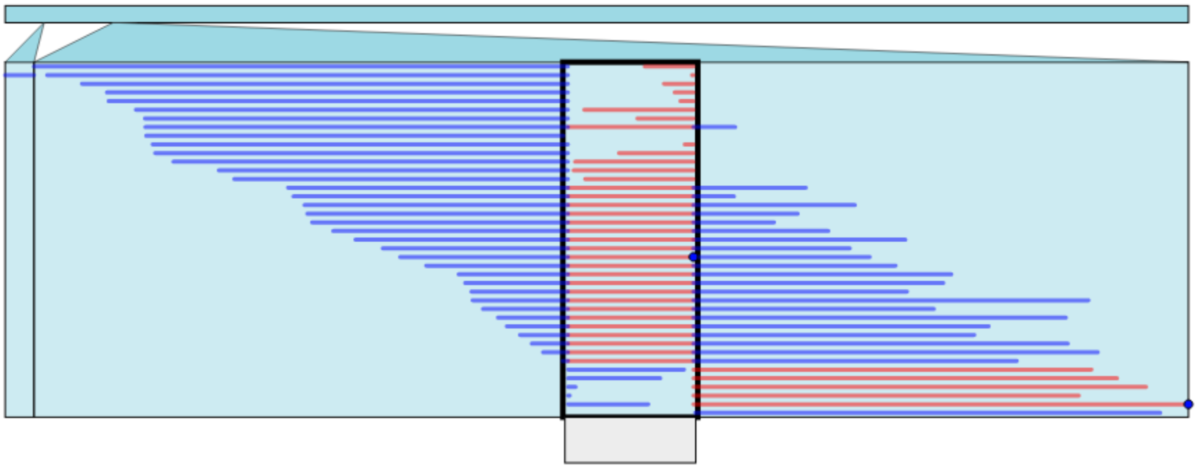

Fig. S19. SM snapshot example: inversion (chromosome_06:1020714-1026818; CC-2931 MA line 15). Ribbon visualisation.

| | | | | | | | | |
|---|---|---|---|---|---|---|---|---|
| F0009 | 2654183 | 17 | 2621271 | - | chromosome_10 | 9087356 | 8766 | 2629767 |
| F0002 | 4417056 | 3409386 | 4417044 | - | chromosome_10 | 9087356 | 2649677 | 3657208 |
| F0002 | 4417056 | 1587364 | 3412473 | - | chromosome_10 | 9087356 | 3656114 | 5481124 |
| F0002 | 4417056 | 776850 | 1587351 | + | chromosome_10 | 9087356 | 5481153 | 6291452 |
| F0002 | 4417056 | 92 | 776832 | - | chromosome_10 | 9087356 | 6291472 | 7068294 |
| F0067 | 483482 | 15 | 483474 | - | chromosome_10 | 9087356 | 7077202 | 7560619 |

Fig. S20. SM snapshot example: inversion (chromosome_10:5481130-6291468; CC-2931 MA line 6). Visualisation from the PAF file. The snapshot includes only a caption of the following fields (i.e. columns): query sequence name (i.e. contig name), query sequence length, query start, query end, relative strand, target sequence name (i.e. chromosome name), target sequence length, target start on original strand, and target end on original strand. Note the lines corresponding to contig F0002 highlighted in white. Genome coordinates in chromosome_10:5481153-6291452 align with a section of contig F0002 that is in reverse orientation compared to the rest of contig sequence, indicating an inversion.

*Translocations*

IGV only enabled visualisation of translocation breakpoints, but reads mapping between ancestral chromosomes were clearly visualised with Ribbon (Fig. S21). All translocations were also inspected from PAF files (e.g. Fig. S22).

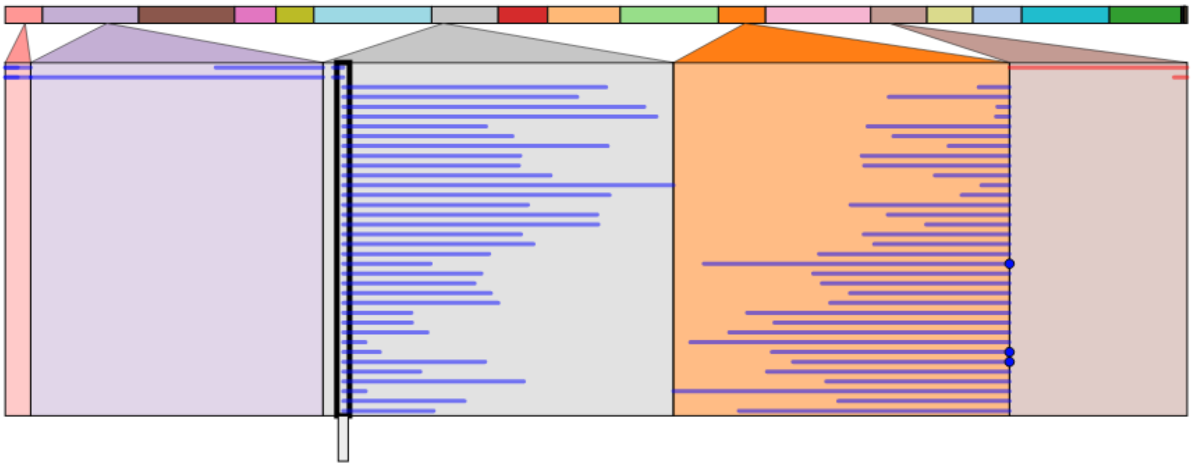

Fig. S21. SM snapshot example: translocation (chromosome_07:1076682 and chromosome_11:2488998; CC-2931 MA line 13). Ribbon visualisation. Notice that the majority of reads map across a breakpoint on the grey-coloured chromosome to the orange coloured chromosome.

| | | | | | | | | |
|---|---|---|---|---|---|---|---|---|
| F0026 | 1652170 | 574420 | 1652164 | - | chromosome 07 | 6159209 | 420 | 1076673 |
| F0007 | 3381888 | 2501012 | 3381880 | + | chromosome_07 | 6159209 | 1076694 | 1957442 |
| F0007 | 3381888 | 6914 | 355782 | + | chromosome_11 | 4356491 | 4 | 348818 |
| F0007 | 3381888 | 1 | 2817 | + | chromosome_11 | 4356491 | 8634 | 11450 |
| F0007 | 3381888 | 350226 | 2500996 | + | chromosome_11 | 4356491 | 338421 | 2488993 |
| F0026 | 1652170 | 304462 | 572594 | - | chromosome_11 | 4356491 | 2488995 | 2757125 |
| F0026 | 1652170 | 11 | 77884 | - | chromosome_11 | 4356491 | 2767239 | 2845118 |
| F0026 | 1652170 | 252042 | 267083 | - | chromosome 11 | 4356491 | 2777329 | 2793002 |

Fig. S22. SM snapshot example: translocation (chromosome_07:1076682 and chromosome_11:2488998; CC-2931 MA line 13). Visualisation from PAF file. The snapshot includes only a caption of the following fields (i.e. columns): query sequence name (i.e. contig name), query sequence length, query start, query end, relative strand, target sequence name (i.e. chromosome name), target sequence length, target start on original strand, and target end on original strand. Note that the lines with a black background correspond to MA line contig F0026, while lines corresponding to contig F0007 are highlighted in white. Note that F0007 transitions from mapping to chromosome 11 (2488993 bp) to mapping to chromosome 7 (1076694 bp). Conversely, F0026 transitions from chromosome 7 to 11 at almost identical coordinates.

### *Transposable element excisions / insertions*

Excisions of transposable elements (TEs) were clearly visualised in IGV. A deletion was often observed together with reads mapping from other copies of the TE elsewhere in the genome (Fig. S23).

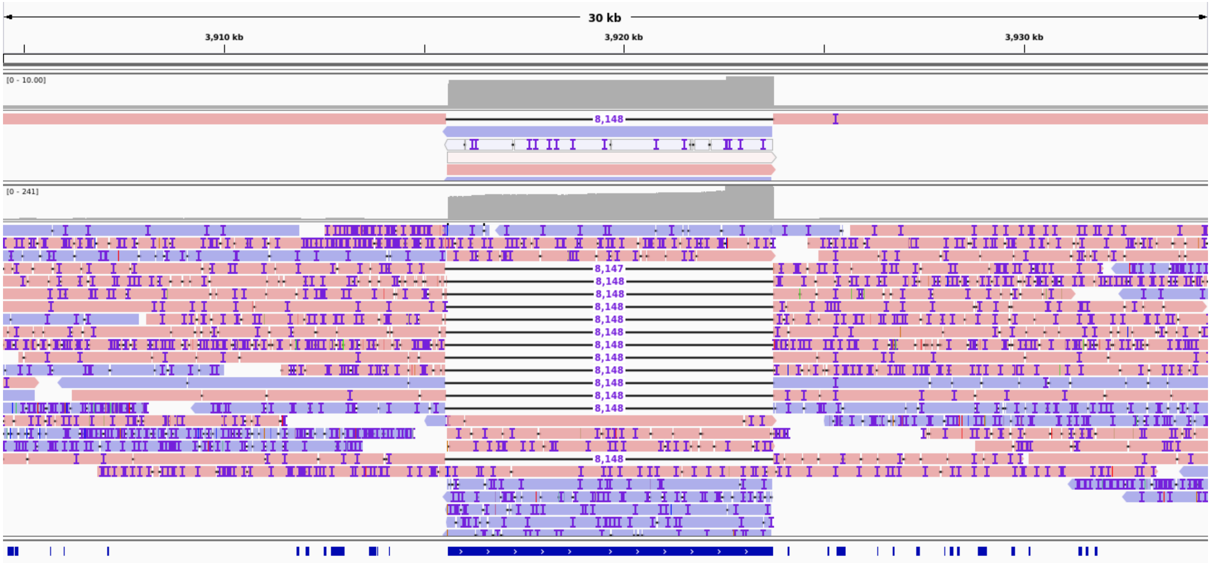


Fig. S23. SM snapshot example: mobile excision (chromosome_16:3915602-3923750; CC-2931 MA line 14). The long blue line at the bottom indicates a TE annotation (*CryptonF-1_cRei*). IGV visualisation. Reads mapping to this region are derived from other copies of this TE present elsewhere in the MA line genome.

TE insertions usually appeared simply as long insertions in the reads, and were clearly visualised in IGV (e.g. an ~8.1 kb *CryptonF-1_cRei* in Fig. S24). Many TE insertions also accompanied breakpoints of other SMs such as inversions and translocations, which were more clearly observed in Ribbon (e.g. a reciprocal translocation mediated by a *U3* element, Fig. S25).

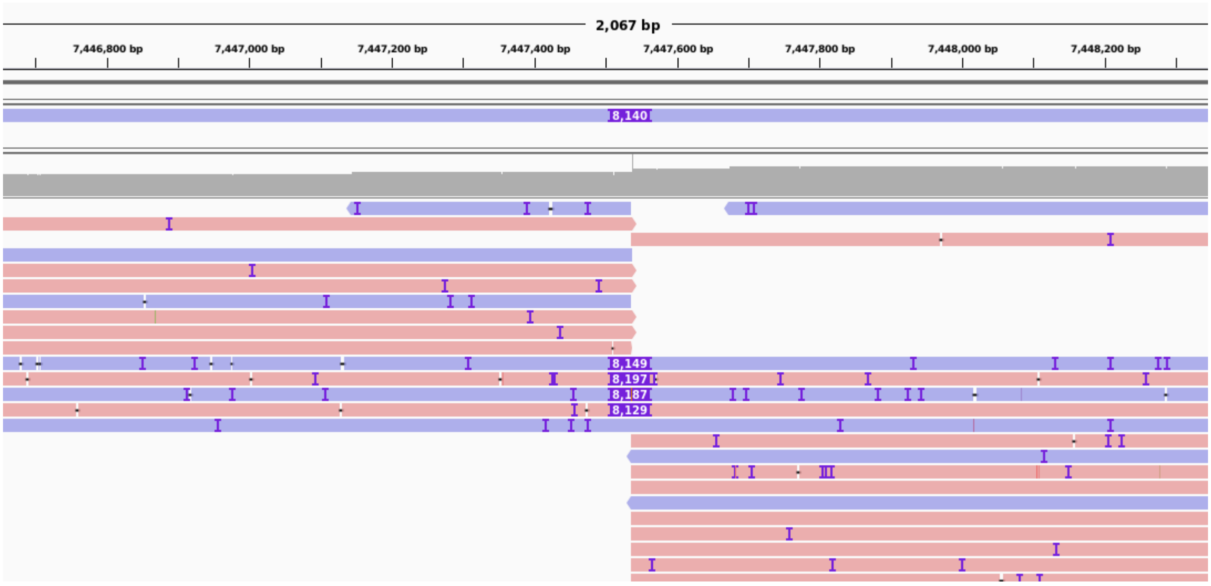


Fig. S24. SM snapshot example: mobile insertion (chromosome_02:7447538-7447539; CC-2931 MA line 14). IGV visualisation.

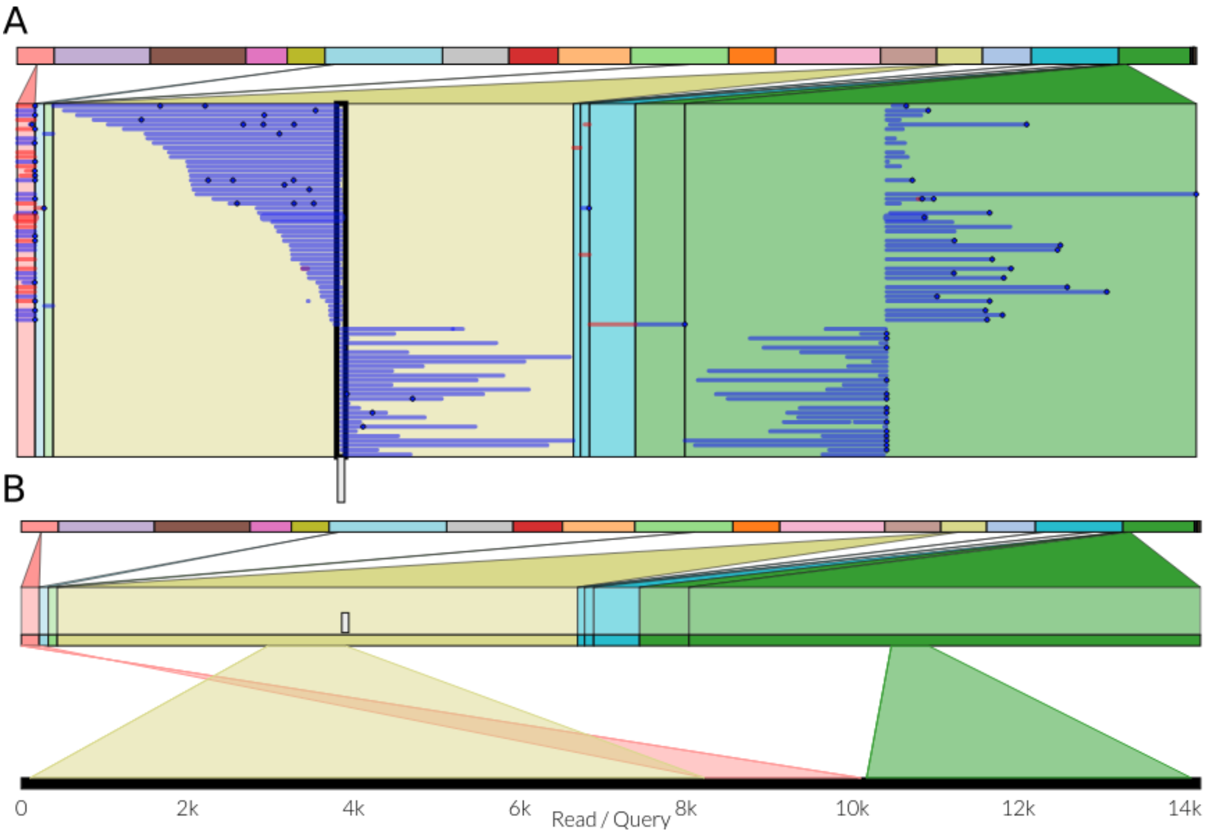


Fig. S25. SM snapshot example: mobile insertion (chromosome_14:1675600; CC-2931 MA line 13) and translocation (chromosome_14:1675600 and chromosome_17:637502). Ribbon visualisation. A) Alignment view. B) Read view (the selected read is highlighted in panel A). The small chromosomal region on the pink chromosome is an ancestral copy of *U3*. The translocation is between the beige and green chromosomes. Note how the reads map from the beige chromosome, through the pink region that contains *U3*, and into the green chromosome. Biologically this signifies a translocation with a *U3* insertion at the breakpoint.

### References


Belyeu JR, Brand H, Wang H, Zhao X, Pedersen BS, Feusier J, Gupta M, Nicholas TJ, Brown J, Baird L et al. 2021. De novo structural mutation rates and gamete-of-origin biases revealed through genome sequencing of 2,396 families. *Am J Hum Genet* **108**: 597-607.

Camacho C, Coulouris G, Avagyan V, Ma N, Papadopoulos J, Bealer K, Madden TL. 2009. BLAST+: architecture and applications. *BMC Bioinformatics* **10**: 421.

Cheng H, Concepcion GT, Feng X, Zhang H, Li H. 2021. Haplotype-resolved de novo assembly using phased assembly graphs with hifiasm. *Nat Methods* **18**: 170-175.

Craig RJ, Hasan AR, Ness RW, Keightley PD. 2021. Comparative genomics of *Chlamydomonas*. *Plant Cell* **33**: 1016-1041.

Gallaher SD, Fitz-Gibbon ST, Strenkert D, Purvine SO, Pellegrini M, Merchant SS. 2018. High-throughput sequencing of the chloroplast and mitochondrion of *Chlamydomonas reinhardtii* to generate improved *de novo* assemblies, analyze expression patterns and transcript speciation, and evaluate diversity among laboratory strains and wild isolates. *Plant J* **93**: 545-565.

Garrison E, Siren J, Novak AM, Hickey G, Eizenga JM, Dawson ET, Jones W, Garg S, Markello C, Lin MF et al. 2018. Variation graph toolkit improves read mapping by representing genetic variation in the reference. *Nat Biotechnol* **36**: 875-879.

Guan D, McCarthy SA, Wood J, Howe K, Wang Y, Durbin R. 2020. Identifying and removing haplotypic duplication in primary genome assemblies. *Bioinformatics* **36**: 2896-2898.

Hunt M, Silva ND, Otto TD, Parkhill J, Keane JA, Harris SR. 2015. Circlator: automated circularization of genome assemblies using long sequencing reads. *Genome Biol* **16**: 294.

Katoh K, Rozewicki J, Yamada KD. 2019. MAFFT online service: multiple sequence alignment, interactive sequence choice and visualization. *Brief Bioinform* **20**: 1160-1166.

Kojima KK, Fujiwara H. 2005. An extraordinary retrotransposon family encoding dual endonucleases. *Genome Res* **15**: 1106-1117.

Kolmogorov M, Yuan J, Lin Y, Pevzner PA. 2019. Assembly of long, error-prone reads using repeat graphs. *Nat Biotechnol* **37**: 540-546.

Koren S, Walenz BP, Berlin K, Miller JR, Bergman NH, Phillippy AM. 2017. Canu: scalable and accurate long-read assembly via adaptive k-mer weighting and repeat separation. *Genome Res* **27**: 722-736.

Larkin MA, Blackshields G, Brown NP, Chenna R, McGettigan PA, McWilliam H, Valentin F, Wallace IM, Wilm A, Lopez R, et al. 2017. Clustal W and Clustal X version 2.0. Bioinformatics 23(1): 2947-2948.

Li H. 2018. Minimap2: pairwise alignment for nucleotide sequences. *Bioinformatics* **34**: 3094-3100.

Manni M, Berkeley MR, Seppey M, Simão FA, Zdobnov EM. 2021. BUSCO update: novel and streamlined workflows along with broader and deeper phylogenetic coverage for scoring of eukaryotic, prokaryotic, and viral genomes. *Mol Biol Evol* doi:10.1093/molbev/msab199.

Nattestad M, Aboukhalil R, Chin CS, Schatz MC. 2021. Ribbon: intuitive visualization for complex genomic variation. *Bioinformatics* **37**: 413-415.

Ness RW, Morgan AD, Vasanthakrishnan RB, Colegrave N, Keightley PD. 2015. Extensive de novo mutation rate variation between individuals and across the genome of *Chlamydomonas reinhardtii*. *Genome Res* **25**: 1739-1749.

Nurk S, Walenz BP, Rhie A, Vollger MR, Logsdon GA, Grothe R, Miga KH, Eichler EE, Phillippy AM, Koren S. 2020. HiCanu: accurate assembly of segmental duplications, satellites, and allelic variants from high-fidelity long reads. *Genome Res* **30**: 1291-1305.

O'Donnell S, Chaux F, Fischer G. 2020. Highly contiguous Nanopore genome assembly of *Chlamydomonas reinhardtii* CC-1690. *Microbiol Resour Announc* **9**: e00726-00720.

O'Donnell S, Fischer G. 2020. MUM&Co: accurate detection of all SV types through whole-genome alignment. *Bioinformatics* **36**: 3242-3243.

Robinson JT, Thorvaldsdóttir H, Winckler W, Guttman M, Lander ES, Getz G, Mesirov JP. 2011. Integrative Genomics Viewer. *Nat Biotechnol* **29**: 24-26.

Ruan J, Li H. 2020. Fast and accurate long-read assembly with wtdbg2. *Nat Methods* **17**: 155-158.

Sedlazeck FJ, Rescheneder P, Smolka M, Fang H, Nattestad M, von Haeseler A, Schatz MC. 2018. Accurate detection of complex structural variations using single-molecule sequencing. *Nat Methods* **15**: 461-468.

Sinha S, Li F, Villarreal D, Shim JH, Yoon S, Myung K, Shim EY, Lee SE. 2017. Microhomology-mediated end joining induces hypermutagenesis at breakpoint junctions. *PLoS Genet* **13**: e1006714.

Smith DR, Lee RW. 2008. Nucleotide diversity in the mitochondrial and nuclear compartments of *Chlamydomonas reinhardtii*: investigating the origins of genome architecture. *BMC Evol Biol* **8**: 156.

Walker BJ, Abeel T, Shea T, Priest M, Abouelliel A, Sakthikumar S, Cuomo CA, Zeng Q, Wortman J, Young SK et al. 2014. Pilon: an integrated tool for comprehensive microbial variant detection and genome assembly improvement. *PLoS One* **9**: e112963.
